# From compartments to gene loops: Functions of the 3D genome in the human brain

**DOI:** 10.1101/2021.10.12.464094

**Authors:** Samir Rahman, Pengfei Dong, Pasha Apontes, Michael B. Fernando, Kayla G. Townsley, Kiran Girdhar, Jaroslav Bendl, Zhiping Shao, Ruth Misir, Nadia Tsankova, Steven P. Kleopoulos, Kristen J. Brennand, John F. Fullard, Panos Roussos

## Abstract

The 3D genome plays a key role in the regulation of gene expression. However, little is known about the spatiotemporal organization of chromatin during human brain development. We investigated the 3D genome in human fetal cortical plate and in adult prefrontal cortical neurons and glia. We found that neurons have weaker compartments than glia that emerge during fetal development. Furthermore, neurons form loop domains whereas glia form compartment domains. We show through CRISPRi on *CNTNAP2* that transcription is coupled to loop domain insulation. Gene regulation during neural development involves increased use of enhancer-promoter and repressor-promoter loops. Finally, transcription is associated with gene loops. Altogether, we provide novel insights into the relationship between gene expression and different scales of chromatin organization in the human brain.

---

The spatiotemporal organization of the genome plays a key role in the regulation of gene expression, the disruption of which has been linked to disease ^1–3^. Chromosomes are partitioned into megabase-sized compartments, named A and B, which contain active and repressed chromatin, respectively ^4^. Compartments are separated into self-interacting neighborhoods called topologically <u>a</u>ssociated <u>d</u>omains (TADs) that are segregated from each other by CTCF bound insulator elements, the disruption of which promotes aberrant enhancer-promoter interactions and transcription mis-regulation ^5–7^. TADs have been proposed to form by loop extrusion of DNA through cohesin, arrested by CTCF bound sites with convergent motifs ^8–11^. Recent work shows dynamic interplay between loop extrusion and phase separation mediated compartmentalization, demonstrating that TADs are not homogeneous structures, but encompass a broad range of features from loop domains to compartment domains, whose respective regulatory functions are poorly understood ^12,13^.

The 3D structure of chromatin changes during neural development ^14^. Studies on mouse and human embryonic stem cells have shown that neural differentiation leads to compartment switching and changes in intra- and inter-TAD interactions ^15,16^. Furthermore, mouse neural differentiation is also associated with the formation of enhancer-promoter loops and *de novo* TADs correlated with neural gene activation ^17^. Highlighting the importance of the 3D genome in disease mechanisms, chromatin interactome studies in both hiPSC derived neurons and human fetal brain revealed that most schizophrenia (SCZ) associated SNPs interact with non-proximal genes that are specifically involved in pathways related to neuronal development and function ^18,19^, corroborating previous work predicting that SCZ associated SNPs disrupt the function of distal enhancer elements ^20^.

Altogether, previous studies have shown changes in 3D genome architecture during neural development; however, little is known about how the hierarchical features of the 3D genome are integrated to regulate gene expression during human brain development. To address this question, we generated high resolution 3D genome maps from human brain samples from different stages of development to elucidate relationships between chromatin architecture and gene expression from the compartment level to intra-genic interactions, highlighting cell type specific differences between neurons and glia and developmental changes across neural development.

## Results

### Neurons disrupt long range B-B compartment interactions and compact A compartments within TADs during development

We performed *in situ* Hi-C on nuclei isolated from *ex vivo* human fetal cortical plate tissue from 18-19 post-conception weeks (pcw) (2 unique donors, defined as early mid gestation fetal) and 23-24 pcw (2 unique donors, defined as late mid gestation fetal), and on neurons and glia from the prefrontal cortex tissue from young (2 unique donors, 30-40 years of age) and old (4 unique donors, 80-100 years of age) adulthood. The 18-24 pcw developmental timeframe represents a period when neurons are completing their migration into the cortical plate ^21^ and extending their synaptic projections ^22–24^, thus transitioning towards functional maturity. After pooling reads from biological replicates we obtained 1.2 billion unique *cis* contacts from fetal cortical plate tissue, and 1.6 and 1.8 billion unique *cis* contacts from adult neurons and glia, respectively, producing ultra-deep 3D genome maps in primary human brain tissue across development (**Supplementary Table 1)**. We measured the reproducibility of Hi-C matrices across biological replicates, and the data clustered according to cell type **(Supplementary Fig. 1a)**. The Hi-C matrices of fetal cortical plate samples correlate more strongly with the matrices of adult neurons than the matrices of adult glia, validating their use to investigate 3D chromatin architecture in developing neurons. Therefore, throughout the manuscript, we refer to fetal cortical plate tissue as “fetal neurons” and neurons isolated from adult brains as “adult neurons”.

We initially examined differences in 3D chromatin structure at the compartment level. We found that fetal and adult neurons have a weaker compartment structure compared to glia (**Fig. 1a, b**). To confirm that the stronger compartment structure in glia is not driven by the heterogeneity of glial cells, we also analyzed the compartment structure in microglia, a specific glial cell type, and found that microglia also have stronger compartments than neurons. We inferred that in contrast to the fine scale compartments in glia, neurons have larger compartments whose borders would overlap less frequently with the boundaries of TADs. Indeed, compared to glia, fetal and adult neurons show less frequent overlap between compartment borders and TAD boundaries (**Fig. 1c**).

**Fig. 1.**
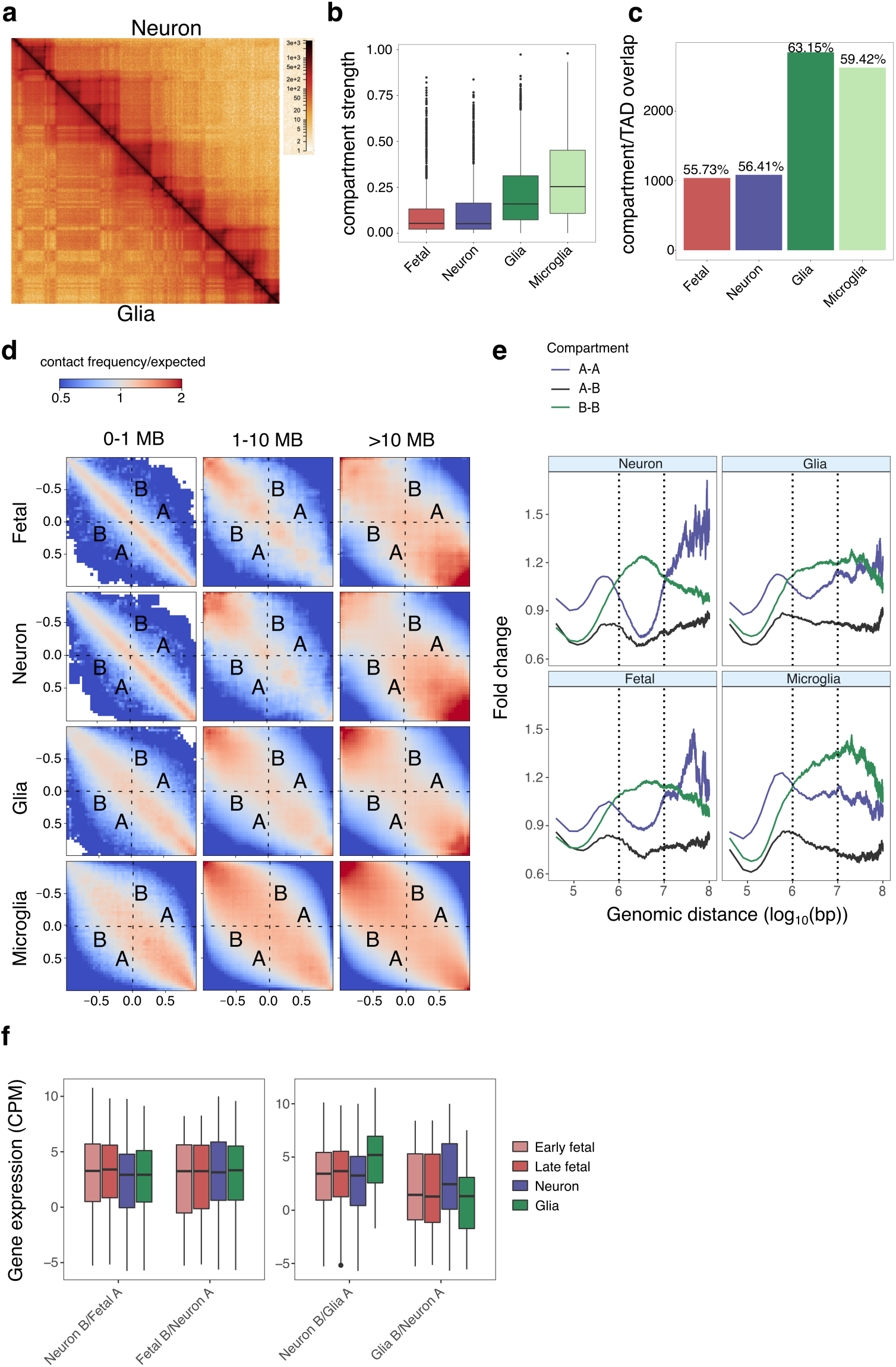
Neurons disrupt long range B-B compartment interactions and compact A compartments within TADs during development. **a**, Snapshot of contact matrices from the HiGlass 3D genome browser representing merged HiC data from adult neurons (upper right) and adult glia (lower left), Chr6:73.1 MB-90.5 MB, at 50 kb resolution. **b**, Compartment strength (determined by CscoreTool) across different cell types. **c**, Frequency of compartment/TAD border overlap across different cell types. Compartment borders within 100 kb of TAD boundaries are considered as overlapping. **d**, Saddle plots generated by cooltools displaying intra-chromosomal interactions in the merged contact maps of fetal neurons, adult neurons, and glia at three genomic distance ranges (0-1 MB, 1-10 MB, and >10 MB) between 40 kb bins ranked by their compartment scores. **e**, Frequency of A-A, A-B, and B-B TAD interactions across genomic distance expressed as fold change above the average interaction frequency of all TADs. **f**, Gene expression (counts per million reads mapped) in loci that switch from compartment B to A between cell types.

As neurons have broader compartments than glia, we hypothesized that the neuronal genome is organized predominantly through loop extrusion, which was shown to disrupt fine scale compartmentalization in recent studies ^12,25^. In particular, we predicted that the intra-chromosomal spatial organization of active (A) and inactive (B) compartments in neurons would be dramatically different compared to glia, as compartmentalization was shown to be induced through B-B attraction ^12^, and loop extrusion was shown to disrupt fine scale B compartments ^25^. Therefore, we investigated the strength of intra-chromosomal A-A and B-B interactions across different ranges of genomic distance (0-1 MB (intra-TAD), 1-10 MB (which we refer to as “inter-TAD” in accordance with a recent study ^26^, and >10 MB (super long-range)), anticipating different patterns of A-A and B-B interactions in neurons compared to glia that are consistent with a loop extrusion model. Interestingly, in adult neurons, we detect a decrease in super long-range B-B compartment interactions when compared to glia (**Fig. 1d**). When we compare A-A interactions between the inter-TAD and intra-TAD ranges, we detect a higher density of intra vs inter TAD A-A interactions in neurons relative to glia, showing that A compartments in neurons have stronger local compaction compared to glia. Fetal neurons show a similar spatial organization of A and B compartments to adult neurons. Conversely, the spatial organization of A and B compartments in microglia is similar to the heterogeneous glial population.

Our observations of stronger intra vs inter-TAD A-A contacts led us to perform a complementary analysis on interactions between TADs, with the hypothesis that neighboring TADs (within the 1-10 MB inter-TAD range) associated with A compartments (i.e. active TADs) would interact less frequently in neurons than in glia. Indeed, we observed that interactions between neighboring inactive TADs (B-B) occur at similar frequencies across all our samples, but interactions between neighboring active TADs (A-A) are much weaker in adult neurons than in glia. (**Fig. 1e**). Altogether, our results show that A and B compartments are organized differently in neurons compared to glia, primarily showing depletion of super long-range B-B compartment interactions, and local compaction of A compartments within TADs.

We then determined the correlation between compartment switching and changes in gene expression. Genes that switch from compartment B to A from neurons to glia are upregulated in glia, whereas genes that switch from compartment B to A from glia to neurons are upregulated in neurons **(Fig. 1f)**. Intriguingly, compartment switching between fetal and adult neurons is associated with relatively modest changes in gene expression in comparison to the differences between adult neurons and glia. Altogether, our findings imply that compartment switching correlates with changes in compartment structure. Given that fetal and adult neurons are very similar in their compartment level structure, most neuron specific A and B compartments may already form by 18 pcw in the cortical plate. As such, developmental gene expression changes due to compartment switching may not be significant.

### Glia form compartment domains of active and inactive chromatin and neurons form loop domains with transcription initiation at boundaries

Most TADs are cell type invariant building blocks of the 3D genome, yet little is known about the cell type specific relationship of TADs to epigenetic modifications and the regulation of transcription. Therefore, we further explored the structure and function of differential TADs in neurons and glia (**Supplementary Fig. 2a**). Having shown that compartment borders and TAD boundaries overlap more frequently in glia compared to neurons, we hypothesized that TADs that are unique to each cell type may be generated through biochemically distinct mechanisms, whereby glia specific TADs may act as compartment domains that form through phase separation of active and inactive chromatin, whereas neuron specific TADs may act as loop domains.

To gain further insight into the mechanisms driving the formation of differential TADs in neurons and glia, we generated ChIP-seq data in the same donors and determined the density of different chromatin states (active enhancer (EnhA) (H3K27ac), active transcription start sites (TssA) (H3K4me3, H3K27ac), and polycomb repressed (ReprPC) (H3K27me3) (**Supplementary Fig. 2b**) across differential TAD boundaries in neurons and glia (**Fig. 2a)**. Compared to cell type invariant TADs, differential TADs in glia show a larger increase in polycomb repression across boundaries relative to neurons, demonstrating that glial TADs have a strong A/B compartment structure. Interestingly, when compared to glia, differential TADs in neurons show stronger enrichment of active TSS at boundaries and downstream active enhancer enrichment. We also detect a stronger compartment shift in glia across glial TAD boundaries relative to neurons, suggesting that while neighboring TADs in glia form their own compartments, those regions converge into a single compartment in neurons (**Fig. 2b)**. Conversely, neuronal TADs have stronger insulation than glial TADs **(Fig. 2c)**. Altogether, differential TADs in glia may form primarily through phase separation of active and inactive chromatin, generating compartment domains, whereas differential TADs in neurons may form primarily through loop extrusion associated with transcriptional activity, explaining the increased insulation at transcription start sites.

**Fig. 2.**
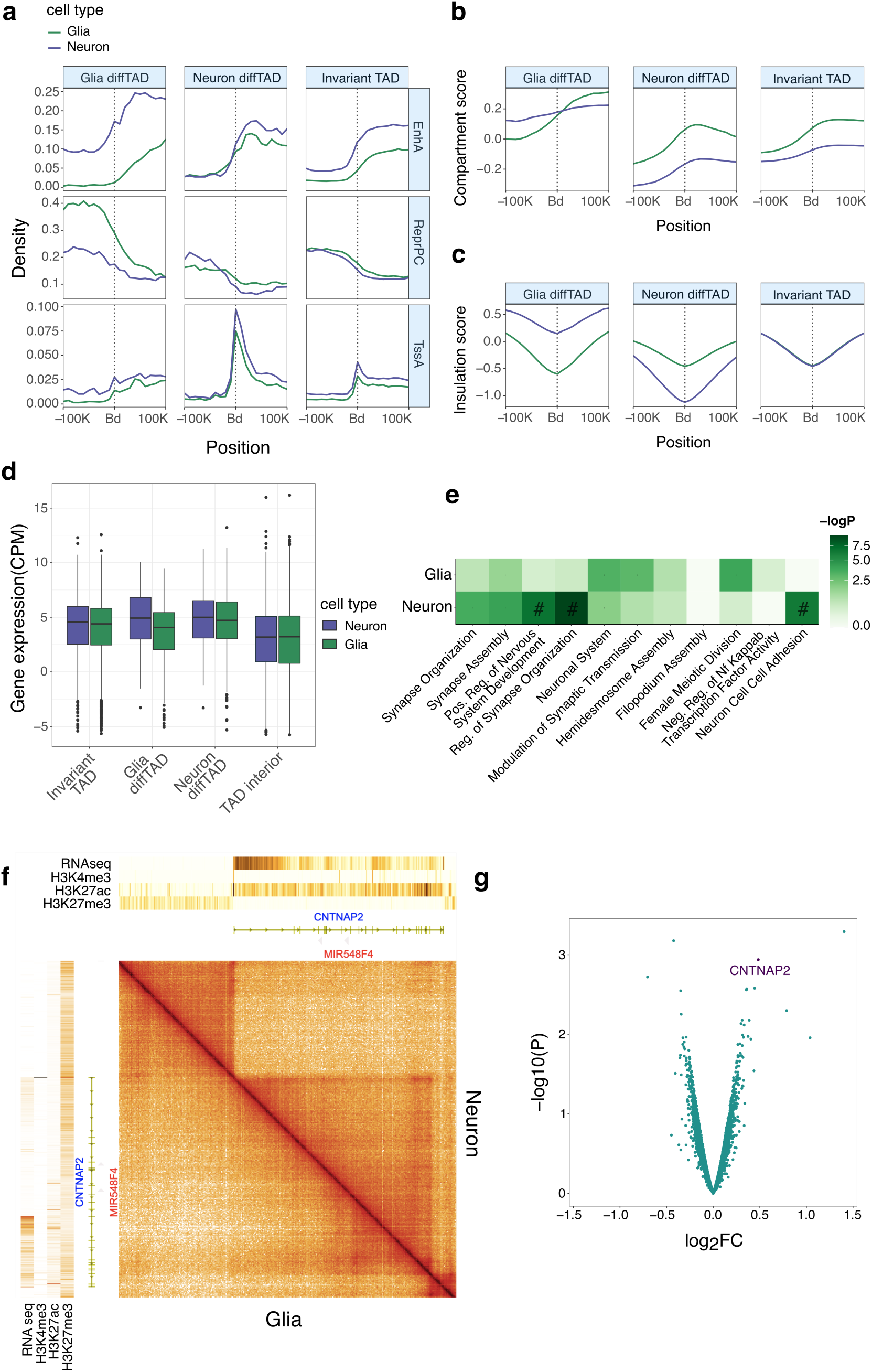
Glia form compartment domains of active and inactive chromatin and neurons form loop domains with transcription initiation at boundaries. **a**, Density of cell type specific chromatin states across differential TAD boundaries (Glia diffTAD: Glia differential TADs, Neuron diffTAD: Neuron differential TADs, Invariant TAD: Cell type invariant TADs). EnhA: active enhancer (H3K27ac), ReprPC: polycomb repressed (H3K27me3), TssA: active transcription start site (H3K4me3, H3K27ac). Bd:Boundary. **b**, Compartment score across differential TAD boundaries. **c**, Insulation score across differential TAD boundaries. **d**, Gene expression (counts per million reads mapped) at differential TAD boundaries. Wilcox p of Invariant=0.001, Wilcox p of Glia=2.95e^−06^, Wilcox p of Neuron=0.0277, Wilcox p of TAD interior=0.0119. **e**, GO terms associated with genes at differential TADs. **f**, Snapshot of the *CNTNAP2* locus from the HiGlass 3D genome browser showing HiC contact maps (10 kb resolution) in neurons and glia along with RNAseq and ChIPseq of H3K4me3, H3K27ac, and H3K27me3 from the respective cell types. **g**, Volcano plot showing the distribution of log2 fold changes in insulation scores between the *CNTNAP2* CRISPRi experiment and scrambled guide control.

We found that differential TAD boundaries in adult neurons are significantly enriched for neuronal gene expression (Fisher’s exact test, odds ratio=1.186, p-value=1.33e^−06^) whereas differential TAD boundaries in glia are depleted of glial gene expression (Fisher’s exact test, odds ratio=0.839, p-value=3.12e^−06^) **(Fig. 2d, Supplementary Fig. 2c)**. Genes at differential TADs in neurons are strongly enriched for processes related to neuronal development **(Fig. 2e)**. Altogether, the regulation of a subset of neuronal genes may be mechanistically coupled to the formation of loop domain boundaries while glial gene regulation may depend primarily on the interior chromatin state of compartment domains.

### *CNTNAP2* forms a loop domain in neurons whose insulation is correlated with ongoing transcription

*CNTNAP2*, the longest gene in the human genome, which is involved in cell-cell adhesion during nervous system development, illustrates the relationships between neuronal differential TAD structure, transcription, and epigenetic regulation. In adult neurons, *CNTNAP2* forms a loop domain initiating from the TSS, showing strong transcription, whereas in glia *CNTNAP2* lacks an active TSS and does not form a loop domain **(Fig. 2f)**. Furthermore, while *CNTNAP2* is enriched in H3K27ac in neurons, in glia it is predominantly enriched in H3K27me3, where it merges with the repressed region upstream. The *CNTNAP2* loop domain is also present in fetal neurons, showing that neuronal TAD structure emerges in the fetal brain and is maintained throughout development (**Supplementary Fig. 2d)**. As glia have higher H3K27me3 enrichment around the 5’ end of *CNTNAP2*, we predicted that this region would be strongly associated with a B compartment in glia whereas in neurons it would tend towards an A compartment. Indeed, the 5’ end of *CNTNAP2* (500 kb downstream from the TSS) is more strongly associated with a B compartment in adult glia and microglia relative to adult neurons (p-value=3.05e^−05^) (**Supplementary Fig. 2e**). Fetal neurons have an intermediate compartment identity between adult neurons and glia at the 5’end of *CNTNAP2*. Altogether, our data suggests that loop domain formation at *CNTNAP2* in neurons is correlated with a focal shift near the TSS from a B compartment towards an A compartment.

To determine if insulation at *CNTNAP2* is coupled to ongoing transcription initiation we performed a CRISPR-interference (CRISPRi) experiment. We recruited dCas9-KRAB to the TSS of *CNTNAP2* in hiPSC derived neurons, resulting in a 20-fold knockdown of *CNTNAP2* expression compared to the scrambled guide negative control experiment (**Supplementary Fig. 2f**). We generated Hi-C data from neurons transduced with the scrambled guide or the guide targeting the *CNTNAP2* TSS and detected a significant and specific decrease in insulation (p-value=0.001) at the *CNTNAP2* TSS in the *CNTNAP2* CRISPRi cells relative to the scrambled control **(Fig. 2g)**. Our result shows that the insulation strength of the *CNTNAP2* loop domain is correlated with ongoing activity at the TSS.

### Neural development involves increased gene regulation through enhancer-promoter and repressor-promoter loops

#### Loops show cell type specific and developmental differences in enrichment of functional chromatin states

We proceeded to investigate chromatin loops to obtain functional insights on regulatory interactions during human brain development. Firstly, we found that loops cluster strongly according to cell type, showing many differences between neurons and glia, but few differences between young and old adult brains **(Fig. 3a)**. Conversely, we find differential loops between early and late fetal neurons, demonstrating that there are more fine scale changes in neuronal 3D genome organization between 18 and 24 pcw, when synaptogenesis is accelerated, than throughout adult brain development. Fetal and adult neurons have longer loops than glia, correlating with published *in vitro* data in which NPC derived neurons were shown to have longer loops compared to NPCs and “astrocyte-like” glia ^19^ (**Supplementary Fig. 3a)**.

**Fig. 3.**
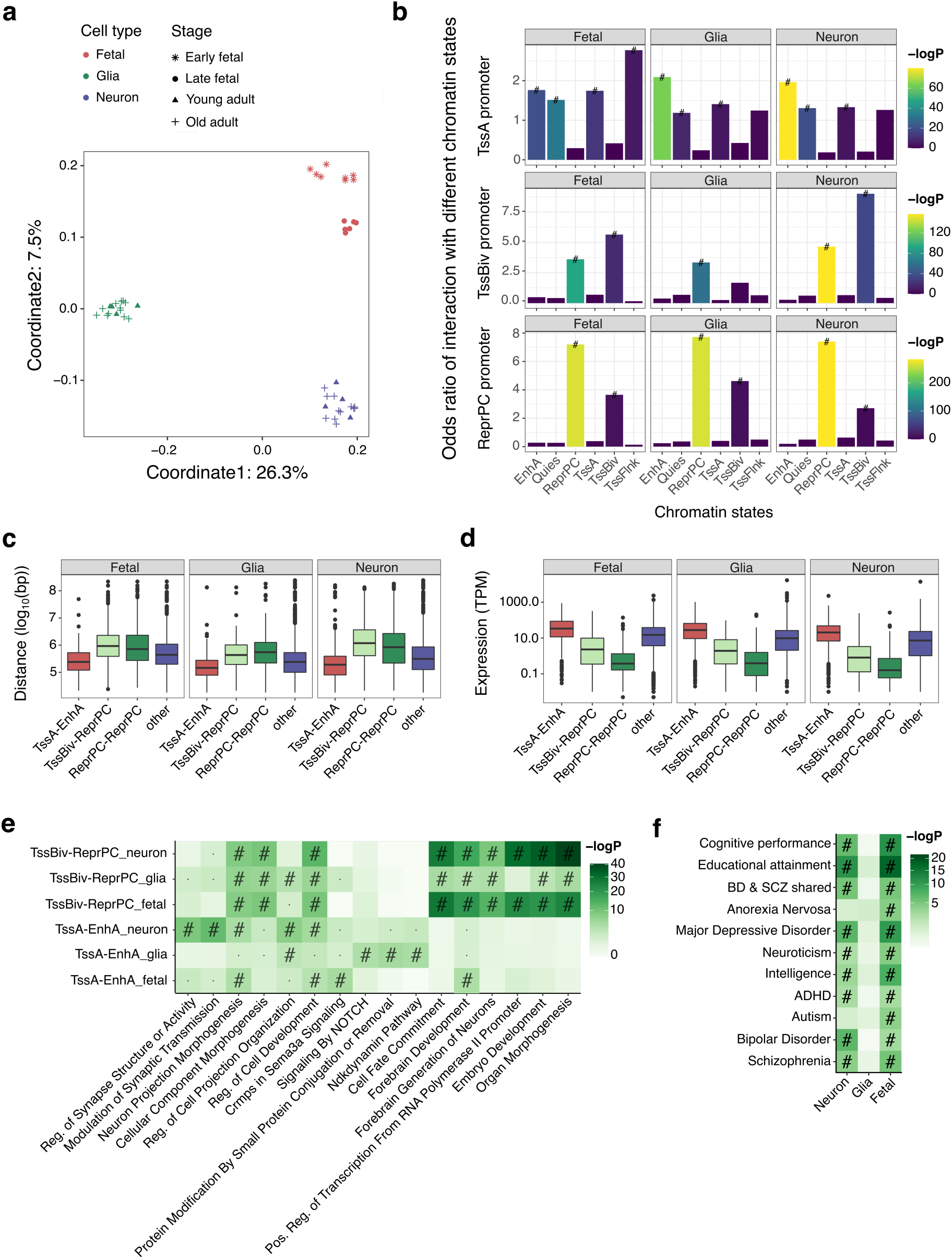
Neural development involves increased gene regulation through enhancer-promoter and repressor-promoter loops. **a**, Multidimensional scaling plot showing differential loops between different cell types and developmental stages (% variance explained). **b**, Odds ratio of interaction between promoter proximal anchors of 3 types (TssA: active transcription start site (H3K4me3, H3K27ac), TssBiv: bivalent transcription start site (H3K4me3, H3K27me3), ReprPC: polycomb repressed (H3K27me3) with chromatin state assigned distal loop anchors. TssFlnk: transcription start site flanking (H3K4me3), EnhA: active enhancer (H3K27ac), Quies: Quiescent. **c**, Distributions of genomic distances (log_10_(bp)) between anchors of TssA-EnhA, TssBiv-ReprPC, and ReprPC-ReprPC loops. **d**, Distributions of expression levels (transcripts per million reads mapped) from genes associated with TssA-EnhA, TssBiv-ReprPC, and ReprPC-ReprPC loops. **e**, GO terms associated with chromatin state assigned differential promoter-anchored loops. **f**, Stratified LD score regression showing the heritability for complex traits and psychiatric disorders within regions associated with differential loops.

We investigated the relationship between the 3D genome and the epigenome in our merged fetal neuron, adult neuron and glia Hi-C contact maps by identifying loops anchored to 3 types of promoters (defined as being within 2kb centered on the TSS): active (TssA) (H3K4me3, H3K27ac), bivalent (TssBiv) (H3K4me3, H3K27me3), and polycomb repressed (ReprPC) (H3K27me3) **(Supplementary Fig. 3b)**, which we predicted could have different levels of activity regulated by distinct distal regulatory elements. We assigned chromatin states to distal loop anchors and found that active promoters in all cell types have a higher probability of interacting with active enhancers (EnhA) relative to other regulatory elements, with a significantly higher probability in adult neurons (**Fig. 3b)**. We found that bivalent and polycomb repressed promoters in all cell types have a higher probability of interacting with polycomb repressed elements, particularly so for adult neurons. Interestingly, bivalent promoters in both fetal and adult neurons have a significantly higher probability of interacting with distal polycomb repressed elements relative to glia. When we compared the proportion of loops that are classified as enhancer-promoter (TssA-EnhA) loops across cell types, we found a higher occurrence of enhancer-promoter loops in adult neurons relative to both fetal neurons (Fisher’s exact test, odds ratio=4.46, p-value= 4.4e^−304^) and glia (Fisher’s exact test, odds ratio=2.5, p-value= 5e^−171^) **(Supplementary Fig. 3c)**. We also found a higher occurrence of TssBiv-ReprPC and ReprPC-ReprPC loops in both fetal and adult neurons relative to glia, with significantly higher frequencies for TssBiv-ReprPC loops (Fisher’s exact test, odds ratio=2.11, p-value= 1.4e^−26^ (Fetal neurons relative to glia), odds ratio=2.45, p-value= 4e^−42^ (Adult neurons relative to glia)). Altogether, our results suggest a cell type specific and age-dependent increase in distal enhancer usage for the regulation of active genes in neurons. In addition, we find a putative role for distal polycomb repressed elements in regulating bivalent promoters throughout neural development.

#### Loops associated with different chromatin states differ in their spatial organization and effect on gene expression

We found that gene expression can be predicted, albeit weakly, from the chromatin state of the distal loop anchor, (Fetal spearman correlation=0.29, Neuron spearman correlation=0.34, Glia spearman correlation=0.31) **(Supplementary Fig. 3d)**. If the distal loop anchor is polycomb repressed, then gene expression is predicted to be downregulated in all cell types. In glia, gene downregulation can also be linked to distal elements that are quiescent, whereas gene downregulation in fetal and adult neurons is exclusively linked to distal polycomb repressed elements. These findings suggest that in glia, genes may be repressed due to the absence of distal activator elements, whereas gene repression during neural development may involve the use of distal elements that have been specifically bookmarked as repressors through increased H3K27me3 density.

We compared the genomic distances between anchors of TssA-EnhA, TssBiv-ReprPC and ReprPC-ReprPC loops and found that polycomb repressed elements tend to interact with target promoters across larger genomic distances than active enhancers (**Fig. 3c)**. We also found that TssA-EnhA loops tend to occur within TADs relative to TssBiv-ReprPC and ReprPC-ReprPC loops (p-value < 2.2e e^−16^), whereas TssBiv-ReprPC and ReprPC-ReprPC loops tend to occur across TADs relative to TssA-EnhA loops (p-value < 2.2e e^−16^) (**Supplementary Table 3a)**. When comparing expression levels across genes associated with TssA-EnhA, TssBiv-ReprPC and ReprPC-ReprPC loops, we find that bivalent promoters are expressed at intermediate levels between active and polycomb repressed promoters (**Fig. 3d)**. Interestingly, when we compare the frequencies of overlap between open chromatin regions obtained from ATAC seq data with our chromatin state assigned loop anchors, we find that the proportion of open chromatin regions that overlap with TssBiv loop anchors is intermediate between the proportions that overlap with TssA and ReprPC loop anchors (**Supplementary Fig. 3e)**. Therefore, promoters interacting with distal polycomb repressed elements may either have highly compacted nucleosomes, leading to strong repression, or partially accessible chromatin states that allow intermediate transcription levels between active and polycomb repressed states. Having shown that bivalent and polycomb repressed promoters in adult neurons have a significantly higher probability of interacting with distal polycomb repressed elements, we find that this stronger tendency to interact with distal polycomb repressed elements correlates with a stronger gene repressive effect. When comparing expression levels between genes associated with differential TssBiv-ReprPC and ReprPC-ReprPC loops across cell types, we find that adult neurons specifically show stronger transcriptional repression at the loop associated promoters relative to fetal neurons and glia (**Supplementary Fig. 3h)**.

Having shown active TSS enrichment at neuronal differential TAD boundaries and active enhancer enrichment downstream of those boundaries, we predicted that there would be more frequent overlap between enhancer-promoter (E-P) loops and differential TADs in neurons relative to glia. Indeed, we found that E-P loops in adult neurons are more likely to be associated with differential TADs relative to glia (Fisher’s exact test, odds ratio=5.51, p-value=1.54×10-^7^), with the promoter anchor more frequently positioned at a differential TAD boundary than the distal enhancer anchor (**Supplementary Table 3b)**. Genes enriched for E-P loops that overlap with neuronal differential TADs are primarily associated with synapse assembly **(Supplementary Table 3c)**. Altogether, a subset of E-P interactions that regulate key neurodevelopmental functions may be facilitated through the formation of loop domains at insulated promoters.

#### Functional significance of differential loops during neural development

We then explored the biological pathways associated with differential loops **(Supplementary Fig. 3i, Fig. 3e)**. TssA-EnhA loops in neurons are enriched for genes primarily associated with glutamatergic neurons, and TssA-EnhA loops in glia are enriched for genes primarily associated with oligodendrocytes, the major non-neuronal cell type in the prefrontal cortex **(Supplementary Fig. 3i)**. TssA-EnhA loops in adult neurons are associated with pathways involved in synaptic structure and activity, while those in fetal neurons are associated with forebrain development **(Fig. 3e)**. Interestingly, TssBiv-ReprPC loops are strongly enriched for developmental processes, implying that polycomb associated repressor elements may fine tune the expression levels of genes that may be critical for spatiotemporal coordination of brain development. Furthermore, only fetal and adult neuron TssBiv-ReprPC loops are enriched for processes related to the positive regulation of transcription from an RNA pol II promoter. Given that neurons are highly specialized cell types, they may need to control RNA Pol II activity to prevent non-specific gene activation. Therefore, polycomb associated repressor elements may downregulate the expression of genes involved in RNA pol II activation.

To obtain insights into neurodevelopmental disease mechanisms we performed stratified LD score regression to determine the heritability for complex traits and psychiatric disorders within differential loops. Interestingly, although both fetal and adult neuron differential loops are significantly associated with SCZ and bipolar disorder (BD), only fetal differential loops are significantly associated with autism **(Fig. 3f)**. Unlike SCZ and BD, whose symptoms emerge during late adolescence, autism emerges during early childhood. Therefore, autism may, in part, be caused by disruptions to regulatory interactions during early brain development.

### Transcription is associated with gene looping and exon-exon interaction strength correlates with isoform expression levels

Brain tissue is biased towards the expression of long genes compared to non-brain tissues, and neurons in particular express longer transcripts compared to non-neuronal brain cell types ^27^. Given the ability of Hi-C to enrich for long range chromatin interactions, we investigated whether transcriptional regulation of long genes in the human brain is associated with gene looping, a phenomenon described in other eukaryotes ^17,28–30^. Firstly, we found that increased gene expression is associated with stronger interactions at TSSs in the direction of transcription **(Fig. 4a)**. We also determined the loop anchor density from the TSS to TES (transcription end site) of each gene **(Fig. 4b)**. Increased gene expression is associated with increased loop anchor density along the gene body, with the highest density found at the TSS. Intriguingly, the loop anchor density decreases from the TSS but increases towards the end of the gene, reaching a peak at the TES and decreasing again downstream of the TES. Furthermore, promoters interact with gene bodies more frequently than downstream regions (**Supplementary Fig. 4c**). Altogether, our results suggest that active promoters scan gene bodies 5’-3’until transcription termination, thus forming dynamic gene loops.

**Fig. 4.**
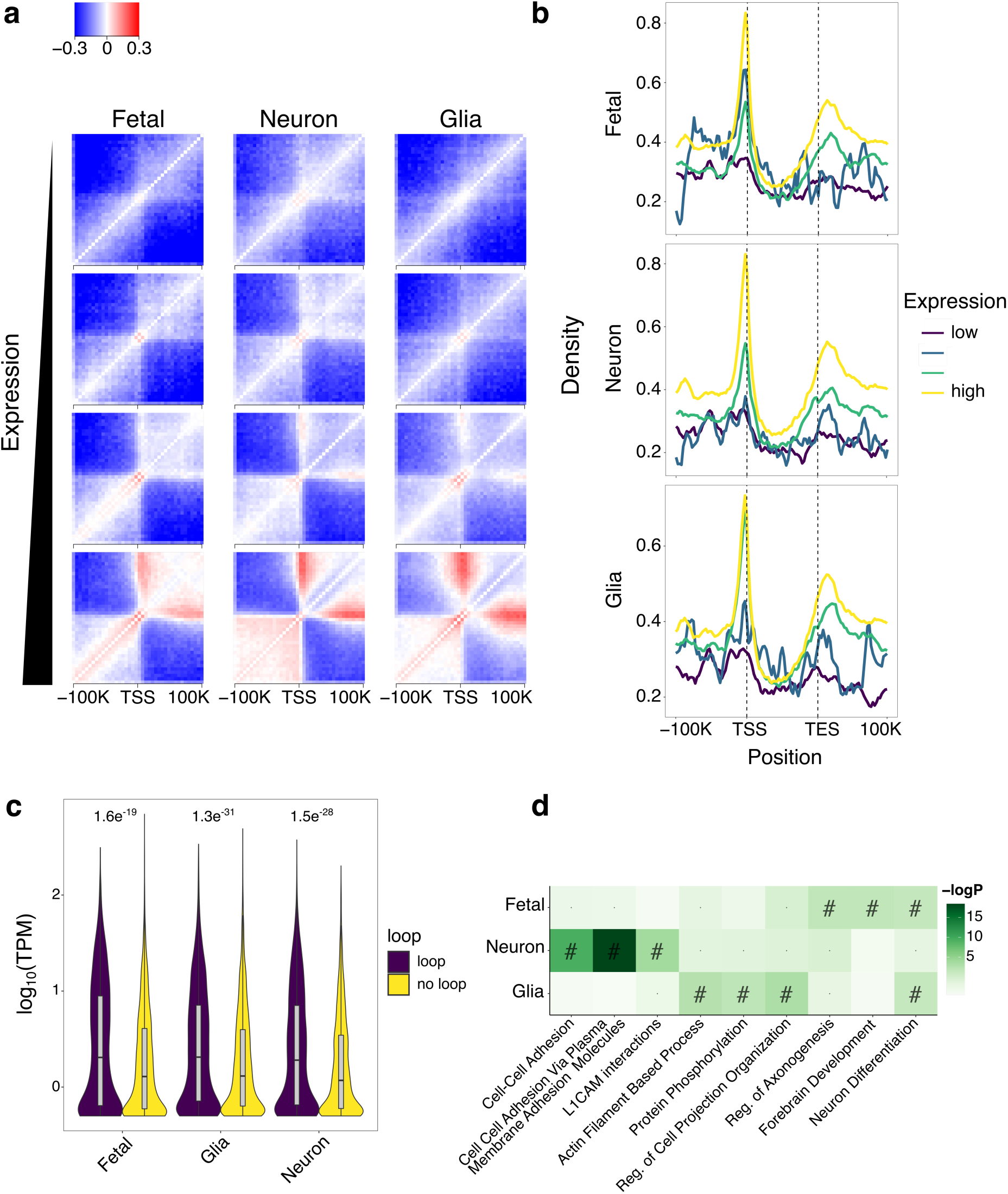
Transcription is associated with gene looping and exon-exon interaction strength correlates with isoform expression levels. **a**, Hi-C interaction heatmap around transcription start sites across 4 different expression levels (TPM: transcripts per million reads mapped) for genes longer than 50 kb. **b**, Loop anchor densities from TSS (transcription start site) to TES (transcription end site) across 4 different expression levels for genes longer than 50 kb. **c**, Violin plots showing fold difference in expression (transcripts per million reads mapped) of isoforms associated with exon-exon loops over isoforms not associated with loops. **d**, GO terms associated with exon-exon loops.

Co-transcriptional splicing is widespread in the human brain ^31^. Intriguingly, a recent study on cohesin-mediated chromatin loops in different cell lines showed a cell type specific increase in the usage of exons that are next to gene body loop anchors linked to the promoter ^32^. These studies led us to hypothesize that intra-gene body chromatin interactions may be involved in co-transcriptional splicing regulation in the human brain. We therefore quantified the relationship between isoform abundance and interactions between exons. Isoforms associated with intra-genic exon-exon loops are more highly expressed than isoforms not associated with exon-exon loops **(Fig. 4c)**. Interestingly, exon-exon loops in fetal neurons are associated with early processes of differentiation, forebrain development, and regulation of axonogenesis, whereas exon-exon loops in adult neurons are associated with processes of cell-cell adhesion that mediate interactions of neurons with their extracellular environment and regulate their excitability **(Fig. 4d)**. These findings correlate 3D gene structure with developmental changes in biological function. Gene folding during transcription may help to coordinate mRNA splicing through exon-exon contacts to generate isoforms with cell type and developmental stage specific functions.

## Discussion

Our work elucidates functional relationships between the 3D genome, the epigenome, and gene expression in the human brain. We focused on large scale differences in compartment and TAD structure between neurons and glia, as well as fine-scale changes in regulatory loops across neural development. We found that neurons have weaker compartments than glia and that this can be detected during fetal development. The weak compartmentalization in neurons could be attributed to the fact that neurons stop dividing as they mature, so they would not replicate DNA, which has been shown to build up compartments and reduce TAD insulation ^33^. The differential intra-chromosomal spatial organization of A and B compartments in neurons relative to glia is suggestive of increased loop extrusion. In particular, the decrease in super long range (> 10 MB) B-B compartment interactions is indicative of strong loop extrusion, as it correlates with findings from a recent study in which depletion of the STAG1 containing variant of cohesin, which is proposed to play a prominent role in TAD stabilization along with CTCF, was shown to increase very long-range interactions between regions that were predominantly located in B compartments ^34^.

The relative enrichment of A-A compartment interactions in intra-TAD (< 1 MB) vs inter-TAD (1-10 MB) genomic intervals in neurons vs glia suggests that loop extrusion in neurons may tend to compact active chromatin locally. Interestingly, a recent study showed that cohesin release by Wapl in mouse embryonic stem cells and neural progenitor cells is required to maintain a cohesin loading cycle to preserve lineage specific transcription ^26^. Wapl depletion leads to decreased intra-TAD (< 1 MB) interactions but increased inter-TAD (1-10 MB) interactions, suggesting that a dynamic cohesin cycle is required to maintain TADs spatially separated from each other. Interestingly, we detect decreased interactions between neighboring active TADs in adult neurons compared to fetal neurons. This finding suggests that the frequency of the cohesin loading cycle may increase during neural development, which, in turn, may increase the probability of neural lineage specific enhancers interacting with their target promoters within TADs rather than forming aberrant interactions across TADs.

We found that differential TADs in neurons and glia, respectively, show different distributions of regulatory histone modifications. Glia specific TADs are characterized by a strong increase in H3K27me3 density across boundaries. As recent work shows that H3K27me3 density is one of the primary drivers of compartmentalization ^35–37^, glial TADs may be considered to be compartment domains that form primarily through polycomb repressive complex mediated phase separation of inactive from active chromatin. Conversely, neuron specific TADs show strong enrichment of active TSSs (H3K4me3, H3K27ac) at boundaries, which are also more insulated than glial TAD boundaries. As cohesin was shown to be recruited to transcription start sites and translocated in the direction of transcription ^38^, along with recent evidence demonstrating that transcription-dependent cohesin repositioning rewires chromatin loops ^39^, neuronal TADs may form primarily through loop extrusion that is driven, at least partially, by transcriptional activity. We performed CRISPRi on *CNTNAP2*, which showed that local transcription inactivation is correlated with specific loss of insulation at the loop domain boundary. Interestingly, a recent study showed that inserting a construct containing a CTCF binding site and a TSS into the genome can generate a *de novo* domain around the insertion site leading to a focal shift from a B compartment towards a more A compartment-like signature ^40^. Our compartment analysis of *CNTNAP2* in different cell types in the human brain correlates with this study, showing that transcriptional activity is coupled to loop domain formation, which may shift chromatin towards an A compartment. Given that fetal neurons have an intermediate compartment identity between adult neurons and glia around the TSS of *CNTNAP2*, this implies that an increase in loop domain strength during neural development correlates with a higher probability of local compartment shifting from B to A.

Interestingly, we find that in comparison to glia a significant subset of E-P loops in neurons overlap with differential TADs and are primarily associated with genes involved in synapse assembly. It’s been shown that promoter loops that are lost upon cohesin degradation have at least one anchor positioned adjacent to a TAD boundary ^41^. Among the E-P loops overlapping with differential TADs, it is the promoter anchor that is more frequently positioned proximally to a differential TAD boundary. Therefore, promoters positioned proximally to differential TAD boundaries in neurons may interact efficiently with their enhancers through loop extrusion, enabling tight enhancer mediated control of gene regulation. Supporting this idea, mouse homologues of some of the genes whose E-P loops overlap with differential TADs (**Supplementary Table 3d)**, including *CNTN5, CNTNAP2, CNTNAP5, FLTR2, FLRT3, GABRA1*, and *GRIA4*, have been shown to be downregulated in mouse cortical neurons upon knockdown of the cohesin component RAD21^42^. As we detect active enhancer enrichment downstream of neuronal TAD boundaries, this may imply a predominantly asymmetric loop extrusion mechanism that reels distal enhancers located downstream towards the promoter. A similar model has been suggested by another study in which CTCF was shown to bind proximally to lineage specific promoters in mouse neural progenitor cells promoting interactions with distal enhancers that are sparsely distributed within the local chromatin environment ^43^. Therefore, loop extrusion may serve as a mechanism to scan local chromatin to facilitate contacts with sparsely populated enhancers. Interestingly, a recent study showed that E-P specificity can be determined by CTCF binding at the promoter, in a cohesin-dependent manner ^44^. Altogether, E-P loops that overlap with differential TADs in neurons may represent a special case of gene regulation in which there is a reciprocal relationship between transcription and 3D structure. In this instance, transcription can facilitate loop extrusion, which allows enhancer scanning, leading to increased transcriptional output from the promoter. Therefore, the recycling of cohesin may be linked to transcriptional memory, which could be important to maintain consistent expression of neurodevelopmental genes. Future work will need to explore the relationships between CTCF, cohesin, neural lineage specific transcription factors, and the general transcriptional machinery in order to elucidate the mechanisms behind the formation of neuronal loop domains.

Our study also finds that repressed and bivalent promoters form loops with distal polycomb repressed elements, providing novel insights on the role of the 3D repressive epigenome in gene supression during neural development. Recent studies show the importance of the polycomb repressive complex 2 (PRC2) in the central nervous system ^45,46^. EZH2, the enzymatic component of PRC2 that deposits H3K27me3, was shown to mediate gene expression patterns in mouse brain that are crucial for neuronal migration ^45^. Also, PRC2 was shown to suppress genes involved in neurodegeneration ^46^. Therefore, long-range polycomb associated loops may play an important role in neuronal survival and function. Polycomb loops may form by phase separation through PRC1, which stabilizes the silencing induced by PRC2, and has the ability to phase separate due to an intrinsically disordered region within its component protein CBX2 ^47^. Therefore, PRC1-PRC1 interactions may bridge polycomb repressed silencer elements with distal promoters, inducing chromatin compaction to ensure suppression of genes, which may play a role in neural development.

Altogether, our survey of the 3D genome during human brain development offers novel insights on the relationship between gene expression and chromatin organization from compartment level structure to gene loops. In particular, we show that cell type specific differences between neurons and glia involve broad scale reorganization of compartment /TAD structure, whereas during neural development to adulthood, changes in the 3D genome involve fine-tuning of regulatory loops.

## Online Methods

### Methods summary

#### Data generation

This study primarily describes Hi-C data generated from nuclei isolated from fetal cortical plate at two stages of development (early mid gestation (18-19 post-conception weeks (pcw)) and late mid gestation (23-24 pcw)) and neuronal (NeuN+) and non-neuronal (NeuN-) nuclei FANS sorted from young adult (30-40 years of age) and old adult (80-100 years of age) prefrontal cortex tissue, using an adaptation of the *in situ* Hi-C protocol ^8^. We generated transcriptome, epigenome, and chromatin accessibility data from the same donors using RNA-seq, Native ChIP-seq (H3K4me3, H2K27ac, and H3K27me3) ^48^, and ATAC-seq ^49^, respectively. We also used complementary Hi-C data generated from microglia that were FACS sorted from fresh human brain samples, as well as Hi-C data generated from iPSC derived neurons, as part of a CRISPRi validation experiment.

#### Data analysis

Hi-C data were aligned with Hi-C pro ^50^, and filtered with hiclib package ^51^ for downstream analysis. The samples are highly reproducible within the same group as determined by HiCRep ^52^. We identified important chromatin organization features including the high-resolution compartments with CscoreTool ^53^, TADs with TopDom ^54^, and chromatin loops with a combination of HICCUPS ^55^ and Fit-Hi-C ^56^. To identify differential loops and TADs, we determined the read counts of loops and the regions between TADs across different samples and used Voom ^57^ and limma ^58^ to normalize the data and determine the significance.

In addition, we have determined chromatin states across different groups with H3K4me3, H3K27ac, and H3K27me3 ChIP-Seq using ChromHMM ^59^. Trait heritability was estimated with Stratified LD score regression ^60^.

### Description of post-mortem brain samples

Fetal brain samples were collected from de-identified prenatal autopsy specimens without neuropathological abnormalities at the Icahn School of Medicine at Mount Sinai (MSSM). The cortical plate was dissected fresh from the anterior frontal lobe of anatomically intact brain specimens. Young and old adult brain samples were accessed through the NIH NeuroBioBanks at Mount Sinai Brain Bank and the University of Miami Brain Endowment Bank. All neuropsychological, diagnostic, and autopsy protocols were approved by the respective Institutional Review Boards.

### Isolation of nuclei from frozen human brain specimens

50 mg of frozen brain tissue was homogenized in chilled lysis buffer (0.32 M Sucrose, 5 mM CaCl_2_, 3 mM Magnesium acetate, 0.1 mM, EDTA, 10 mM Tris-HCl, pH 8, 1 mM DTT, 0.1% Triton X-100) and filtered through a 40 µm cell strainer. Filtered lysate was then underlaid with sucrose solution (1.8 M Sucrose, 3 mM Mg(CH_3_COO)_2_, 1 mM DTT, 10 mM Tris-HCl, pH 8) and subjected to ultracentrifugation at 107,000 xg for 1 hour at 4 °C. Pellets were resuspended in 500 µl DPBS containing 0.1% BSA. For non-fetal tissue, anti-NeuN antibody (1:1000, Alexa488 conjugated, Millipore, Cat# MAB377X) was added and samples incubated, in the dark, for 1 hr at 4 °C. Prior to FANS sorting, DAPI (Thermoscientific) was added to a final concentration of 1 µg/ml. DAPI positive neuronal (NeuN+) and non-neuronal (NeuN-) nuclei were isolated using a FACSAria flow cytometer with FACSDiva Version 8.0.1 software (BD Biosciences). In the case of fetal tissue, where NeuN is not expressed, nuclei were isolated using DAPI staining only.

### Isolation of microglia from fresh human brain specimens

Fresh autopsy and biopsy tissue specimens were processed using the Adult Brain Dissociation Kit (Miltenyi Biotech Cat# 130107677), according to manufacturer’s instructions. Following de-myelination (Miltenyi de-myelination kit, Miltenyi Biotech, Cat# 130096733) cells were incubated in antibody (CD45: BD Pharmingen, Clone HI30 and CD11b: BD Pharmingen, Clone ICRF44) at 1:500 for 1 hour in the dark at 4 °C with end-over-end rotation. RNase inhibitors (Takara Bio) were used throughout the cell prep. Prior to fluorescence activated cell sorting (FACS), DAPI (Thermoscientific) was added at 1:1000 to facilitate the separation of live from dead cells. Viable (DAPI negative) CD45/CD11b positive cells were isolated by FACS using a FACSAria flow cytometer (BD Biosciences). Following FACS, cell concentrations and viability were confirmed using a Countess automated cell counter (Life technologies).

### *in situ* Hi-C on frozen human brain tissue

Hi-C data was generated from frozen postmortem human brain tissue using the *in situ* Hi-C protocol ^8^ with the following modifications. Frozen prefrontal cortex tissue was thawed at room temperature (RT) and dounce homogenized in HBSS (Hank’s balanced salt solution). Homogenized tissue was fixed with 0.5% formaldehyde for 10 min and then quenched with 0.125 M glycine for 5 min at RT. Cross-linked tissue was then placed on ice for a further 15 min to quench crosslinking completely. Tissue was pelleted at 800 xg for 10 min at 4 °C and then resuspended in lysis buffer (0.32 M Sucrose, 5 mM CaCl_2_, 3 mM Mg(CH_3_COO)_2_, 0.1 mM EDTA, 10 mM Tris-HCl, pH 8, 1 mM DTT, 0.1 % Triton X-100, 1 Roche cOmplete mini EDTA-free protease inhibitor tablet, Roche Cat# 4693159001) to isolate cross-linked nuclei.

Approximately 1 M crosslinked neuronal and non-neuronal (glia) nuclei from adult brains, or nuclei from fetal cortical plate, were thawed on ice, washed with ice cold 1x CutSmart buffer (New England Biolabs (NEB), Cat# B7204S) and split into 4 aliquots to generate technical replicate libraries per sample, with 250k nuclei per library. Nuclei were pelleted at 2500 xg for 5 min at 4 °C, and resuspended in 342 µl 1x CutSmart buffer and then conditioned with 0.1% SDS at 65 °C for 10 min. Nuclei were immediately placed on ice and the SDS quenched with 1% Triton X-100. Chromatin was digested with 100 U of the 4 base pair cutter MboI (New England Biolabs, Cat# R0147L) overnight at 37 °C with shaking at 400 rpm. MboI was heat inactivated at 65 °C for 20 min, and then nuclei were cooled down on ice. MboI cut sites were end labeled with biotin by adding 52 µl of biotin fill-in reaction mix (15 µl of 1 mM biotin-14-dATP (Jena Bioscience, Cat# NU-835-BIO14-L), 1.5 µl each of 10 mM dCTP, dGTP, dTTP (Sigma-Aldrich, Cat# DNTP10-1KT), 10 µl of 5 U/µl Klenow DNA Pol I (New England Biolabs, Cat# M0210L), 22.5 µl of 1x CutSmart buffer) and shaking at 37 °C for 1.5 h at 400 rpm. Blunt ended sites were proximity ligated by adding 948 µl ligation reaction mix (150 µl of 10x T4 DNA ligase buffer (New England Biolabs, Cat# B0202S), 125 µl of 10% Triton X-100, 15 µl of 10 mg/ml BSA, 10 µl of 400 U/µl T4 DNA ligase (New England Biolabs, Cat# M0202L), 648 µl ddH_2_0) and rotating tubes, end-over-end, at RT for 4 h. Nuclei were reverse crosslinked with 100 µl proteinase K (10 mg/ml) overnight at 65 °C.

Proximity ligated DNA was purified through phenol:chloroform extraction and sodium acetate/ethanol precipitation. Purified DNA was sheared using a Covaris S220 sonicator to generate a peak size of 400 bp with the following settings (peak incident power: 140 W, duty cycle: 10%, cycles per burst: 200, time: 55 sec). Biotin labeled ligation junctions were purified with Dynabeads MyOne Streptavidin C1 beads (ThermoFisher, Cat# 65001) by incubating for 1 hr at RT. Illumina compatible libraries were prepared from the sonicated and streptavidin bead immobilized DNA using the NEBNext Ultra II Library prep kit (New England Biolabs, Cat# E7645L), following manufacturer’s instructions, amplifying libraries for 6-13 PCR cycles. Libraries were purified by 2-sided size selection (300-800 bp) using Ampure XP beads (Beckman Coulter, Cat# A63881). All libraries were analyzed on a TapeStation using Agilent D5000 ScreenTapes, and quantified using the KAPA Library Quantification Kit prior to sequencing. Uniquely barcoded Hi-C libraries were pooled and deep sequenced on the Illumina NovaSeq S4 platform obtaining 100 bp paired end reads, to generate approximately 250 million reads per library.

### *in situ* Hi-C on microglia sorted from fresh human brain tissue

Hi-C libraries from microglia were generated using a modified version of the *in situ* Hi-C protocol ^8^ or the Arima HiC+ kit.

For the *in situ* Hi-C protocol, 500k-1 M FACS sorted microglia were fixed with 1% formaldehyde for 10 min at RT and then quenched with 0.125 M Glycine for 5 min at RT. Crosslinking was quenched completely by incubating cells on ice for 15 min. Cells were pelleted at 800xg for 5 min at 4 C, washed in 1xPBS. Cells were either frozen on dry ice and stored at −80 C or processed for the experiment immediately.

Fixed cells were resuspended in the *in situ* Hi-C lysis buffer (10 mM Tris-HCl (pH=8.0), 10 mM NaCl, 0.2% Igepal CA-630 (NP40), 1 Roche cOmplete mini EDTA-free protease inhibitor tablet, Roche Cat# 4693159001). Cells were lysed for 15 min on ice. Nuclei were pelleted at 2500xg for 5 min at 4 C and then washed with ice cold 1x CutSmart buffer (New England Biolabs, Cat# B7204S) and split into 4 aliquots to generate technical replicate libraries per sample, with about 200k nuclei per library. Nuclei were pelleted at 2500xg for 5 min at 4 °C, and resuspended in 342 µl 1x CutSmart buffer and then conditioned with 0.1% SDS at 65 °C for 10 min. Nuclei were immediately placed on ice and the SDS quenched with 1% Triton X-100. Chromatin was digested with 100 U of the 4 base pair cutter MboI (New England Biolabs, Cat# R0147L) overnight at 37 °C with shaking at 400 rpm. MboI was heat inactivated at 65 °C for 20 min, and then nuclei were cooled down on ice. MboI cut sites were end labeled with biotin by adding 52 µl of biotin fill-in reaction mix (15 µl of 1 mM biotin-14-dATP (Jena Bioscience, Cat# NU-835-BIO14-L), 1.5 µl each of 10 mM dCTP, dGTP, dTTP (Sigma-Aldrich, Cat# DNTP10-1KT), 10 µl of 5 U/µl Klenow DNA Pol I (New England Biolabs, Cat# M0210L), 22.5 µl of 1x CutSmart buffer) and shaking at 37 °C for 1.5 h at 400 rpm. Blunt ended sites were proximity ligated by adding 948 µl ligation reaction mix (150 µl of 10x T4 DNA ligase buffer (New England Biolabs, Cat# B0202S), 125 µl of 10% Triton X-100, 15 µl of 10 mg/ml BSA, 10 µl of 400 U/µl T4 DNA ligase (New England Biolabs, Cat# M0202L), 648 µl ddH_2_0) and rotating tubes, end-over-end, at RT for 4 h. Nuclei were reverse crosslinked with 100 µl proteinase K (10 mg/ml) overnight at 65 °C.

Proximity ligated DNA was purified through phenol: chloroform extraction and sodium acetate/ethanol precipitation. Purified DNA was sheared using a Covaris S220 sonicator to generate a peak size of 400 bp with the following settings (peak incident power: 140 W, duty cycle: 10%, cycles per burst: 200, time: 55 sec). Biotin labeled ligation junctions were purified with Dynabeads MyOne Streptavidin C1 beads (ThermoFisher, Cat# 65001) by incubating for 1 hr at RT. Illumina compatible libraries were prepared from the sonicated and streptavidin bead immobilized DNA using the NEBNext Ultra II Library prep kit (New England Biolabs, Cat# E7645L), following manufacturer’s instructions, amplifying libraries for 8-10 PCR cycles. Libraries were purified by 2-sided size selection (300-800 bp) using Ampure XP beads (Beckman Coulter, Cat# A63881). All libraries were analyzed on a TapeStation using Agilent D5000 ScreenTapes, and quantified using the KAPA Library Quantification Kit prior to sequencing. Uniquely barcoded Hi-C libraries were pooled and deep sequenced on the Illumina NovaSeq S4 platform obtaining 100 bp paired end reads, to generate approximately 250 million reads per library.

For the Arima HiC protocol, microglia were fixed with 2% formaldehyde for 10 min at RT and then quenched with 0.125 M Glycine for 5 min at RT. Crosslinking was quenched completely by incubating cells on ice for 15 min. Cells were pelleted at 800xg for 5 min at 4 C, washed in 1xPBS, and then split into 4 aliquots to generate technical replicate libraries per sample. Hi-C was performed according to the manufacturer’s instructions. Proximity ligated DNA was purified with Ampure XP beads and then Hi-C libraries were prepared in identical fashion as the *in situ* Hi-C protocol.

### Native ChIP-seq

Ultra-low input Native Chromatin Immunoprecipitation sequencing (ULI-NChIP-seq) libraries were prepared as follows. After FANs-sorting of neuronal (NeuN+) and non-neuronal nuclei (NeuN-nuclei), we performed ULI-NChIP assays, adapted from ^48^, which specifically does not require chromatin crosslinking, thereby increasing library complexity and reducing PCR artifacts. Briefly, nuclei were centrifuged at 500xg for 10 minutes at 4 °C and re-suspended by gentle pipetting in residual smaller volumes of PBS/Sheath buffer. After DAPI counting of nuclei, 100K-300K were distributed into Eppendorf tubes and 0.1% Triton-X-100/0.1% Na-Deoxycholate added. The chromatin was re-suspended and placed at room temperature for 5 minutes, followed by fragmentation with micrococcal nuclease (MNase, NEB, Cat#M0247S) for 5 min at 37 °C on a ThermoMixer at 800 rpm to digest the chromatin to predominantly mononucleosomes. The MNase reaction was stopped by addition of 10% of the reaction volume of 100 mM EDTA (pipetted ~20x), followed by the addition of 1% Triton / 1% deoxycholate, (pipetted 5x) and placed on ice for at least 15 minutes. Samples were vortexed (medium setting) for ~ 30 seconds and complete NChIP buffer (20 mM Tris-HCl, pH 8.0, 2 mM EDTA, 15 mM NaCl, 0.1% Triton X-100, 1 EDTA-free protease inhibitor cocktail and 1 mM phenylmethanesulfonyl fluoride) was added to dilute the chromatin to <25% of the immunoprecipitation reaction volume, and rotated at 4 °C for 1 hour. After the incubation, chromatin was vortexed (medium setting) for ~30 seconds and 5% input controls removed for DNA extraction.

To avoid non-specific binding, chromatin was next precleared by adding 10 µl/ reaction of a pre-washed 1:1 ratio of protein A to protein G Dynabeads (ThermoFisher, Cat# 10001D and #10003D), rotating the chromatin-protein A/ protein G magnetic bead mixture for 3 hours at 4 °C. Antibody-bead complexes were prepared as follows: ChIP-grade histone antibodies H3K27ac (Active Motif, Cat# 39133, pAB), H3K4me3 (Cell Signaling, Cat# 9751), or H3K27me3 (Cell Signaling, Cat# 9733) were added to prewashed protein A/protein G Dynabeads resuspended in NChIP buffer, and the antibody-bead complexes formed by rotating the antibody and beads for 2 hours at 4 °C. After the incubations, precleared chromatin and antibody-bead complexes were placed on a magnetic rack and the precleared chromatin transferred to new Eppendorf tubes, while the antibody-bead complexes were resuspended in sufficient volume of NChIP buffer to add 10 µl per MNase reaction. The chromatin was immunoprecipitated with the antibody-bead complexes at 4 °C overnight while rotating.

After the overnight incubation, the immunoprecipitation reactions were placed on a magnetic rack to remove unbound chromatin, washed twice with 200 µl of ChIP low salt wash buffer (20 mM Tris-HCl, pH 8.0, 0.1% SDS, 1% Triton X-100, 0.1% deoxycholate, 2 mM EDTA and 150 mM NaCl), twice with 200 µl ChIP high salt wash buffer (20 mM Tris-HCl (pH 8.0), 0.1% SDS, 1% Triton X-100, 0.1% deoxycholate, 2 mM EDTA and 500 mM NaCl), followed by elution in freshly prepared 30 µl of ChIP elution (100 mM sodium bicarbonate and 1% SDS) for 1.5 hours at 68°C on a ThermoMixer at 1000 rpm, RNAse A digestion for 15 minutes at 37 °C at 800 rpm. The immunoprecipitated DNA, along with the input controls, was purified using Phenol:Chloroform:Isoamyl Alcohol (25:24:1, v/v) (ThermoFisher Scientific, Cat# 15593-031), transferred to pre-spun phase lock tubes (Qiagen Maxtract, Cat# 129046) to obtain the aqueous layer. An overnight ethanol precipitation was performed by adding to the 10 µl of 3M sodium acetate/100 µl aqueous layer of ChIP DNA, 1 µl of LPA (linear polyacrylamide, Sigma, Cat# 56575) and 1 µl Glycoblue (Invitrogen, Cat#AM9515).

NEBNext Ultra I DNA Library Prep Kit (New England Biolabs, Cat# E7370L) was used to construct Native ChIP-seq libraries according to the manufacturer’s directions, followed by Pippin Size selection using 2% Agarose Gel cassettes (SAGE Science) and cleanup with 1.8 volumes of SPRIselect beads (Beckman Coulter, #B23318). All libraries were analyzed on a TapeStation using Agilent High Sensitivity D1000 ScreenTapes, and quantified using the KAPA Library Quantification Kit prior to sequencing. The uniquely barcoded libraries were pooled and sequenced on the Illumina NovaSeq S4 platform obtaining 100 bp paired end reads. Approximately 40 million paired-end reads were generated per sample and subsequently aligned on hg38.

### RNA-Seq

For RNA-seq, nuclei were sorted into 1.5 ml low-binding Eppendorf tubes containing Extraction buffer, a component of the PicoPure RNA Extraction kit (Arcturus, Cat# KIT0204). RNA was isolated in accordance with the PicoPure RNA Isolation kit’s manufacturer’s instructions. This included an RNase-free DNase treatment step (Qiagen, Cat # 79254), which was completed according to the instructions provided by the manufacturer. Samples were eluted in RNase-free water and stored at −80 °C until preparation of RNA-Sequencing libraries using the SMARTer Stranded Total RNA-Seq Pico Kit v1 or v2 (Takara Clontech Laboratories, Cat# 635005 or 634414, respectively), according to the manufacturer’s instructions. Following construction of the RNA-seq libraries, libraries were analyzed on a TapeStation using a High Sensitivity D1000 ScreenTape (Agilent, Cat# 5067-5584) and quantification of the libraries was performed using the KAPA Library Quantification Kit. RNA-seq libraries were subsequently sequenced on the Illumina NovaSeq S4 platform yielding 100 bp paired-end reads.

### ATAC-Seq

ATAC-seq libraries were generated using an established protocol^49^. Briefly, 55,000 to 75,000 sorted nuclei were pelleted at 500 xg for 10 min at 4 °C. Pellets were resuspended in transposase reaction mix (22.5 µL Nuclease Free H_2_O, 25 µL 2x TD Buffer; Illumina, Cat # FC-121-1030) and 2.5 µL Tn5 Transposase (Illumina, Cat # FC-121-1030) on ice and the reactions incubated at 37 °C for 30 min. Following incubation, samples were purified using the MinElute Reaction Cleanup kit (Qiagen Cat# 28204, and libraries generated using the Nextera index kit; Illumina Cat #FC-121-1011). Following amplification, libraries were resolved on 2% agarose gels and fragments ranging in size from 100-1000 bp were excised and purified (Qiagen Minelute Gel Extraction Kit– Qiagen, Cat# 28604). Next, libraries were quantified by quantitative PCR (KAPA Biosystems, Cat# KK4873) and library fragment sizes estimated using Tapestation D5000 ScreenTapes (Agilent technologies, Cat# 5067-5588). ATAC-seq libraries were subsequently sequenced on the Illumina NovaSeq S4 platform yielding 100 bp paired-end reads.

### CRISPRi guide design and plasmid preparation

The CRISPRi algorithm of the Broad Institute Genetic Perturbation Platform was used to design guides against *CNTNAP2*. The top 3 guides picked by the algorithm were synthesized by IDT (Integrated DNA technologies) and cloned downstream of a constitutively expressed U6 promoter in a lentiviral vector (lentiGuide-Hygro-mTagBFP2, Addgene, Cat# 99374) using the golden gate cloning method. Guide oligos and their reverse complement oligos were initially phosphorylated and annealed using the following protocol:

1 μl guide oligo (100 μM)

1 μl reverse complement oligo (100 μM)

1 μl 10X T4 DNA Ligase buffer (New England Biolabs, Cat# B0202S)

0.5 μl T4 PNK (New England Biolabs, Cat# M0201L)

<u>6.5 μl ddH2O</u>

10 μl total

Thermocycler:

37 °C 30 min

95 °C 5 min

Ramp down to 25 °C at 5 °C /min

4 °C infinite hold

Phospho-annealed oligos were diluted 1:100 before proceeding to the golden gate reaction:

12.5 μl Quick Ligation Buffer (2X) (New England Biolabs, Cat# M2200L)

0.25 μl BSA (10 mg/ml)

1 μl BsmB1 v2 (New England Biolabs, Cat# R0739L) (optimized for golden gate cloning)

0.125 μl T7 DNA Ligase (New England Biolabs, Cat# M0318S)

1 μl diluted phospho-annealed oligos

1 μl of [25ng/ul] LV-EF1a-U6 backbone

<u>9.125 μl ddH2O</u>

25 μl total

Thermocycler:

Cycle (30x):

37 °C 5min

20 °C 5min

4 °C infinite hold

Ligations were transformed into High Efficiency NEB 10-beta Competent *E. coli*. (New England Biolabs, Cat# C3019H) using the manufacturer’s protocol. Bacteria were plated on LB-Ampicillin agar medium at 37 °C overnight. On the following day, colonies were grown in LB-Ampicillin liquid medium at 37 °C with shaking. Plasmids were purified with the QIAprep Spin Miniprep kit (Qiagen, Cat#27106). Positive clones were validated through sanger sequencing by GeneWiz using a universal U6 forward primer. Vectors were packaged into viruses by VectorBuilder.

### NPC CRISPRi

hiPSCs and NPCs (NSB553-S1-1 and NSB2607-1-4) were derived by sendai viral OKSM reprogramming of dermal fibroblasts obtained from control donors and differentiated using dual-SMAD inhibition as described previously^61^. Both donors are male and of European ancestry.

Established CRISPRi dCas9-KRAB colonized control hiPSC-NPC lines^62^ (NSB553-S1-1 and NSB2607-1-4), were maintained in Matrigel (Corning, Cat# 354230) coated 6-well plates under NPC medium (DMEM/F12 (Life Technologies, Cat# 10565), 1x N2 (Life Technologies, Cat# 17502-048), 1x B27-RA (Life Technologies, Cat# 12587-010), 20 ng/ml FGF2 (R&D Systems, Cat# 233-FB-01M). Upon confluency, cells were dissociated with Accutase (Innovative Cell Technologies, Cat# AT104) for 5 minutes at 37 °C, quenched with DMEM/F12, pelleted and resuspended in NPC medium containing 10uM/ml Thiazovivin (THX) (Sigma/Millipore, Cat# 420220). 3.5×10^5 NPCs per well were seeded onto Matrigel coated 24-well plates in NPC media. On the following day, gRNA lentiviruses such as LentiGuide-Hygro-mTagBFP2 (Addgene, Cat# 99374) were added to cultures, followed by spinfection (1 hour, 1000xg, 25°C). After spinfection, the cultures were incubated overnight, and medium was then replaced the following day. The cells were selected with 0.3 µg/ml puromycin for dCas9-KRAB (Sigma, Cat# P7255) for two days. After puromycin selection, cells were fed with fresh NPC medium and then harvested two days later. gRNA expression was confirmed via BFP fluorescence, prior to harvest.

### Neuron induction

At day-2, hiPSC-NPCs were seeded as 5-8×10^5^ cells / well in a 24-well plate coated with Matrigel or 1.0-1.2×10^6^ cell/well in a 12-well plate <u>with THX</u>. At day-1, cells were transduced with rtTA (Addgene, Cat#20342), *NGN2-Neo or NGN2-Neo-GFP* (Addgene, Cat# 99378) and gRNA (Addgene, Cat# 99374) lentiviruses via spinfection. At Day 0, 1 µg/ml dox was added to induce *NGN2*-expression. At Day 1, transduced hiPSC-NPCs were treated with corresponding antibiotics to the lentiviruses (0.3ug/ml puromycin for dCas9-KRAB-Puro, 0.5 mg/ml G-418 for *NGN2*-Neo and 0.400 mg/ml HygroB for lentiguide-Hygro-mTagBFP2) in order to increase the purity of transduced hiPSC-NPCs. At day 3, NPC medium was switched to neuronal medium (Brainphys (Stemcell Technologies, Cat# 05790), 1x N2 (Life Technologies, Cat# 17502-048), 1x B27-RA (Life Technologies Cat# 12587-010), 1 µg/ml Natural Mouse Laminin (Life Technologies), 20 ng/ml BDNF (Peprotech, Cat #450-02), 20 ng/ml GDNF (Peptrotech, Cat# 450-10), 500 µg/ml Dibutyryl cyclic-AMP (Sigma, Cat# D0627), 200 nM L-ascorbic acid (Sigma, Cat# A0278)) including 1 µg/ml Dox, along with antibiotic withdrawal. 50% of the medium was replaced with fresh neuronal medium. At day 11, full medium change withdrew residual dox completely. At day 13, *NGN2*-excitatory neurons were treated with 200 nM Ara-C to reduce the proliferation of non-neuronal cells in the culture, followed by half medium change by day 17. At Day 17, Ara-C was completely withdrawn by full medium change, followed by half medium changes until the neurons were fixed or harvested day 21-24.

### CRISPRi validation

RNA was extracted from transduced Ngn2 neurons (NSB2607-1-4) using the QIAzol lysis reagent (Qiagen, Cat# 79306) and purified using the Qiagen RNAeasy MinElute Cleanup Kit (Qiagen, Cat# 74204). RNA was quantified using the Qubit RNA HS Assay Kit (ThermoFisher, Cat# Q32852) and RNA integrity was determined using the Agilent High Sensitivity RNA ScreenTape. cDNA was generated using the High Capacity cDNA Reverse Transcription Kit (ThermoFisher, Cat# 4368813), following the manufacturer’s protocol using an input of 400 ng RNA per cDNA synthesis reaction. Target gene expression knockdown was validated through qPCR using TaqMan probes (ThermoFisher, Cat# 4331182 and 4351370) and the TaqMan Gene Expression Master Mix (ThermoFisher, Cat# 4369510). qPCR was performed in triplicate for every sample using the manufacturer’s protocol. The 2^−∆∆Ct^ method was used to determine fold change in target gene expression between the knockdown experiment and the scrambled guide control relative to the housekeeping genes GAPDH and ACTB.

### *in situ* Hi-C on CRISPRi validated Ngn2 induced neurons

Transduced Ngn2 neurons were grown in 12 well plates for *in situ* Hi-C. Media was removed and cells were washed once with 1x PBS, and then fixed with 2% formaldehyde/1x PBS for 10 min at RT. Formaldehyde fixation was quenched with 0.125 M glycine for 5 min at RT. Plates were then kept on ice for 15 min to quench cross-linking completely. Cells were gently scraped off the wells into 1.5 ml eppendorf tubes. Cells were pelleted at 800xg for 5 min at 4 °C and then washed twice with 1x PBS. After removing the supernatant, cells were frozen on dry ice for 20 min and then stored at −80 °C.

Hi-C libraries were prepared from cells (NSB2607-1-4) that were transduced with the top scoring guide for *CNTNAP2* (replicate libraries from 3 wells), which was shown by TaqMan qPCR to downregulate gene expression at least 20-fold (**Fig. S2F**), and cells (NSB2607-1-4) that were transduced with the scrambled guide (replicate libraries from 6 wells). Libraries were prepared using the Arima-HiC+ kit, following manufacturer’s instructions, with the following modifications. Purified proximity ligated DNA was sheared using a Covaris S220 sonicator to generate a peak size of 400 bp with the following settings (peak incident power: 140 W, duty cycle: 10%, cycles per burst: 200, time: 55 sec). Biotin labeled ligation junctions were purified with Dynabeads MyOne Streptavidin C1 beads (ThermoFisher, Cat# 65001) by incubating for 1 hr at RT. Illumina compatible libraries were prepared from the sonicated and streptavidin bead immobilized DNA using the NEBNext Ultra II Library prep kit (NEB, Cat# E7645L), following manufacturer’s instructions, amplifying libraries for 6-11 PCR cycles. Libraries were purified by 2-sided size selection (300-800 bp) using Ampure XP beads (Beckman Coulter, Cat# A63881). All libraries were analyzed on a TapeStation using Agilent D5000 ScreenTapes, and quantified using the KAPA Library Quantification Kit prior to sequencing. Uniquely barcoded Hi-C libraries were pooled and deep sequenced on the Illumina NovaSeq S4 platform obtaining 100 bp paired end reads, to generate approximately 250 million reads per library.

### RNA-seq data processing

The trimmed reads were aligned to human genome hg38 (GRCh38) using STAR (2.7.2a) aligner^63^, where the allelic alignment bias was corrected by WASP^64^. Gene expression was quantified using RSEM (v1.3.1) tools^65^ with GENCODE V30 as a reference and summarized at both gene and isoform levels. Duplication and GC content levels were estimated by Picard tools (v2.2.4). Quality control (QC) metrics were collected by RNA-seq QC (v1.1.8)^66^.

### ATAC-seq data processing

#### Alignment

The trimmed reads were aligned to human genome hg38 (GRCh38) using STAR aligner^63^ with specific parameters for ATAC-Seq alignment (“--alignIntronMax 1 -- outFilterMismatchNmax 100 --outFilterScoreMinOverLread 0.66 -- outFilterMatchNminOverLread 0.66 -alignEndsType Local”), and controlled allelic alignment bias with WASP^64^. Reads mapped to multiple loci detected by samtools (v0.1.19), duplicated reads that were marked by Picard (v2.2.4), and mitochondria alignments were filtered.

#### Peak calling

We called peaks for glia, neuron, and fetal independently. For each cell type, bam files from each sample were subsampled to the same sequencing depth and then merged. We used model-based Analysis of ChIP-seq (MACS, v2.1)^67^ to call peaks with a smoothing window of 200bps (--shift -100 --extsize 200 --nomodel) and FDR of 0.01 (-q 0.01). Peaks overlapped with ENCODE blacklist regions^68^ were discarded, yielding 222,746 neuronal, 136,171 glia, and 149,412 in fetal.

### ChIP-seq data processing

H3K4me3, H3K27ac, H3K27me3 ChIP-seq, and corresponding input files were aligned to human genome hg38, similar to our ATAC-seq pipeline, above. The resulting bam files were subsampled to the same sequencing depth and merged for each cell type. We called peaks using MACS with a smoothing window of 150bps (--shift -75 --extsize 150 --nomodel), FDR of 0.01 (-q 0.01), and input as control. For H3K27me3, we called broad peaks (--broad). Peaks overlapped with ENCODE blacklist regions^68^ were discarded.

### Chromatin states annotation

We implemented a multivariate Hidden Markov Model model (ChromHMM)^59^ to systematically annotate the combinational effect of different histone modifications. The ChromHMM model is trained by virtually concatenating histone marks H3K4me3, H3K27ac, and H3K27me3 in all three cell types that merged across all individuals that subsampled to uniform depths. Reads were shifted from 5’ to 3’ direction by 100 bp for all the samples. Read counts were then computed in 200bp non-overlapping bins across the genome. Each bin was binarized into 1 or 0 by the Poisson model with a p-value threshold of 10^−4^. We have trained the model with merged data using six states which captured all the key interactions from our data. Lastly, we obtained the chromatin states with the trained model and corresponding binarized files as input.

### Hi-C data processing

#### Alignment and filtering

Hi-C data were aligned using the HiC-Pro strategy^50^. Briefly, paired-end reads were mapped independently to the human genome hg38 using bowtie2 in stringent mode with parameters (“--very-sensitive -L 20 --score-min L,-0.6,-0.2 --end-to-end”)^69^. Then, the chimeric reads that failed to align were trimmed after ligation sites (MboI “GATCGATC”) and mapped to the genome. All the aligned reads from both ends were then merged based on read names and mapped to MboI restriction fragments using hiclib package^51^. Next, self-circles, dangling ends, PCR duplicates, and genome assembly errors were discarded. Samples of the same cell type were merged. We binned the interaction matrix at different resolutions and corrected it with iterative correction (ICE) for downstream analysis. We collected Hi-C specific quality control information for each technical replicate, including 1) percent of input chimeric reads. 2) alignment summary from HiC-Pro. 3) quality control from hiclib package, including self-circle, dangling ends, PCR duplicates, percent of cis/trans reads. 4) slop, the slope of the linear regression between log scaled average interaction frequency and log scaled average genomic distance (100Kb-10Mb, 100Kb resolution). 5) percent of contacts within between TADs (insulation region) (For TAD analysis).

### Similarities between replicates

To determine the reproducibility across different replicates of our Hi-C data, we used HiCRep package^52^. HiCRep stratified the interaction matrix by genomic distance and then determined the stratum adjusted correlation coefficient (SCC) to compare the similarities. We compared the biological replicates of neuron, fetal, and glia samples at 100kb resolution. The samples exhibit strong within-group correlation.

### High-resolution compartment analysis

To annotate high-resolution A/B compartments for each of the different tissue and stage, we used a two-step strategy. First, we performed principal component analysis using a genome-wide interaction matrix at 200kb. We compared the correlation between the resulting first principal component (PC1) and gene density of each bin, and reverse the sign if it’s a negative correlation. Thus, we generated a genome-wide compartment score at low resolution. Next, for each chromosome, we used CscoreTool^53^ to annotate A/B compartments at 40kb bin resolution. We reversed the sign of the Cscore if it correlates negatively with the genome-wide PC1 score. We discard the chromosomes where the PC1 and Cscore have a low correlation (Pearson correlation

<0.5). In this way, we generated a high-resolution compartment annotation for each group. To identify compartment switch, we convert the difference between two tissues or stages into z-scores and get the corresponding p-value small than 0.01 bins as compartment switch loci.We also analyzed the strength of the compartment by generating saddle plots using the cooltools package (https://github.com/open2c/cooltools/tree/master/cooltools). Briefly, intra-chromosome interactions were normalized and averaged for each genomic distance at 40kb bin. Then the distance normalized interactions were aggregated by the high-resolution compartment score. In addition, we have assigned TAD into A/B compartment based on the compartment coverage. Then we normalized the A-A, A-B, and B-B TAD interactions for each genomic distance (Figure 1E).

### TAD analysis

Topological associated domains (TADs) were identified with Topdom^54^ at 10K resolution and a 200Kb window size.

In order to quantify the dynamic of TADs during the development process as well as after CRISPR interference, we determined the insulation score^70^ of all the shared TAD borders across different technical replicates. Read counts were then normalized with the trimmed mean of M-values (TMM) method^71^. We performed a PCA on the normalized read count matrix for each assay to identify high-variance components that explained at least 1% of the variance. Correlation tests were performed between the selected PCs and known covariates, and covariates with FDR<0.05 were used for the following steps. To select the final covariates, we first chose the covariates such as cell types and sex that are known to play a critical role for each assay as “a base model”. We then applied an approach based on the Bayesian information criterion (BIC) to select the final covariates^72^. We examined the BIC changes in the linear regression model after adding a new covariate, which will be included if it can improve the mean BIC by at least 4. In different comparisons across development, we used covariates including person ID, Sex, Pool, slope, percent of cis-reads, and percent of contacts between TADs (insulation region). For CRISPR interference analysis, we used covariates including slope, percent of trans reads, percent of dangling ends, PCR duplicates, non-unique map reads.

With the selected covariates, the normalized read counts were modeled with the voomWithQualityWeights function from the limma package (v.3.38.3)^58^, which utilizes both sample-level and observational-level weights. We subsequently perform the test against the contrast between tissues or between CRISPR treatment using a linear mixed model to account for repeated measurements (i.e. 2 brain regions per individual) in the dream function^73^ of the variancePartition package^74^.

In order to annotate the differential TAD borders, we determined the ChromHMM annotation, compartment score, and insulation score across the 100kb window up/downstream of the differential TAD borders at 10kb bins. The side with a lower compartment score was set as upstream (−100k). We also collected functional gene sets from MSigDB 7.0^75^, and used one-tailed Fisher exact tests to test the enrichment and significance of the differential TAD borders associated genes.

### Chromatin loop analysis

Chromatin loops were identified with both HICCUPS^55^ and Fit-Hi-C^56^ of merged Hi-C contacts for each tissue^76^. For Fit-Hi-C, loops were called from 5k to 20k (by 1k) resolution. In order to get a consistent P cutoff, we used the P cutoff of FDR (Benjamini-Hochberg adjusted) <0.05 at 10kb for different resolutions. After that, we get the chromatin loops that are reproducible in at least to different resolutions. For HICCUPS, chromatin loops were called using juicer HICCUPS with bin sizes iterated from 10kb to 25kb by 1kb intervals and parameters “-k VC_SQRT -p 1 -i 3”. Only reproducible loops were retained and the highest resolution of the overlapping loops was used. Lastly, loops were merged from the two methods.

In order to annotate differential loops, we determined the interaction counts at 20kb resolution across all the technical replicates. Then we used the same pipeline as the differential TAD analysis to identify differential chromatin loops.

In order to annotate all the promoter-associated loops, we get all the promoters (1kb of TSS) that within the chromatin loop anchors. We get the chromatin states at the promoter loci, as well as the chromatin states at the distal region. Then we assign the promoter and the distal region to the chromatin state that has the largest coverage. We fit a 10-fold cross-validation lasso model with the glmnet package (v 2.0.18)^77^, use distal chromatin states, or both promoter and distal chromatin states to predict the gene expression.

We collected functional gene sets from MSigDB 7.0^75^, and human brain single-cell markers^78^ and perform one-tailed Fisher exact tests to test the enrichment and significance of loop associated genes.

### Partitioned heritability analysis

We partitioned heritability for loop anchors to examine the enrichment of common variants in neuropsychiatric traits with stratified LD score regression (v.1.0.0)^60^ from a selection of GWAS studies^79–85^. Briefly, with the loop anchors, a binary annotation was created by marking all HapMap3 SNPs^86^ that fell within the loop anchors and outside the MHC regions. LD scores were calculated for the overlapped SNPs using an LD window of 1cM using 1000 Genomes European Phase LD reference panel^87^. The enrichment was determined against the baseline model^60^.

## Acknowledgments

We thank the patients and families who donated material for these studies. We thank the computational resources and staff expertise provided by the Scientific Computing group at the Icahn School of Medicine at Mount Sinai.

## Funding

Supported by the National Institute on Aging, NIH grants U01-MH116442 (to P.R.), R01-MH109897 (to P.R. and K.J.B.), R01-MH109677 (to P.R.) and R01-MH110921 (to P.R.). R01MH106056 (to K.J.B), U01DA047880 (to K.J.B), R01DA048279 (to K.J.B), 6R56MH101454 (to K.J.B), and R01MH121074 (to K.J.B.). J.B. is partially supported by a NARSAD Young Investigator Grant 27209 from the Brain and Behavior Research Foundation (BBRF). P.D. is partially supported by a NARSAD Young Investigator Grant 29683 from the Brain and Behavior Research Foundation (BBRF). K.G. is partially supported by an Alzheimer’s Association Research Fellowship (AARF-21-722582).

## Author contributions

S.R., P.D. and P.R. conceived of and initiated the project. S.R., J.F.F. and P.R. designed experimental strategies for omics profiling. N.T. provided human fetal brain tissue. S.R., P.A., Z.S., S.P.K., R.M., and J.F.F. performed omics data generation in human brain tissue. M.B.F, K.G.T., S.R. and K.J.B. performed the CRISPRi *in vitro* validation studies. S.R., P.D. and P.R. designed analytical strategies. P.D., K.G. and J.B. conducted initial bioinformatics, sample processing and quality control for the omics data. P.D. developed the computational scheme and performed the downstream analysis. S.R., J.F.F. and P.R. supervised overall data generation. S.R., P.D. and P.R. supervised overall data analysis. S.R., P.D. and P.R. wrote the manuscript with input from all authors.

## Competing interests

The authors declare no competing interests.

## Supplementary Figures

**Supplementary Fig. 1.**
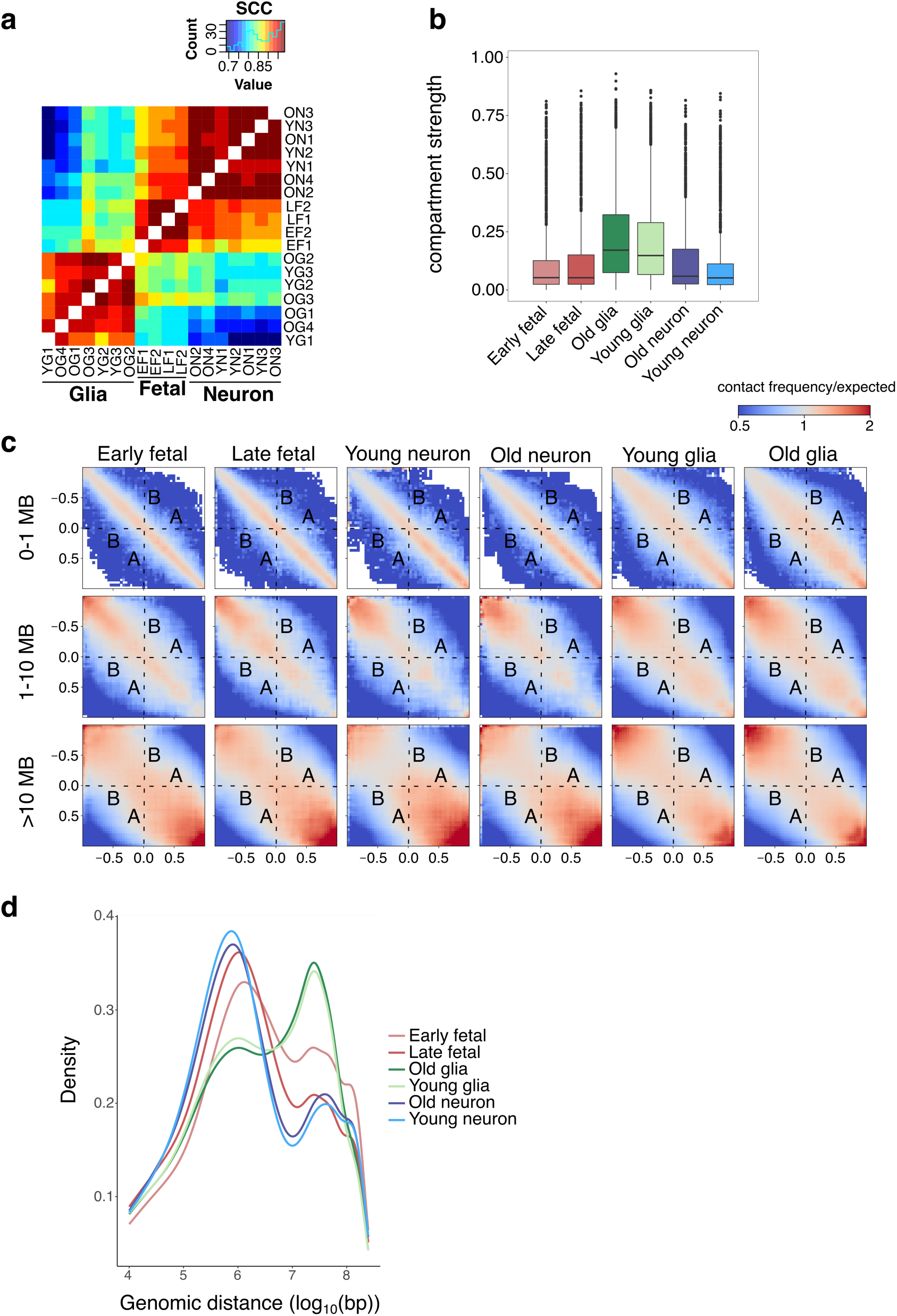
**a**, Stratum adjusted correlation coefficients (SCC) between Hi-C matrices from biological replicates determined by HiCRep. **b**, Compartment strength (determined by CscoreTool) across different cell types and developmental stages. **c**, Saddle plots generated by cooltools displaying intra-chromosomal interactions at three genomic distance ranges (0-1 MB, 1-10 MB, and >10 MB) between 40 kb bins ranked by their compartment scores. **d**, Interaction frequencies across genomic distance (log_10_(bp)).

**Supplementary Fig. 2.**
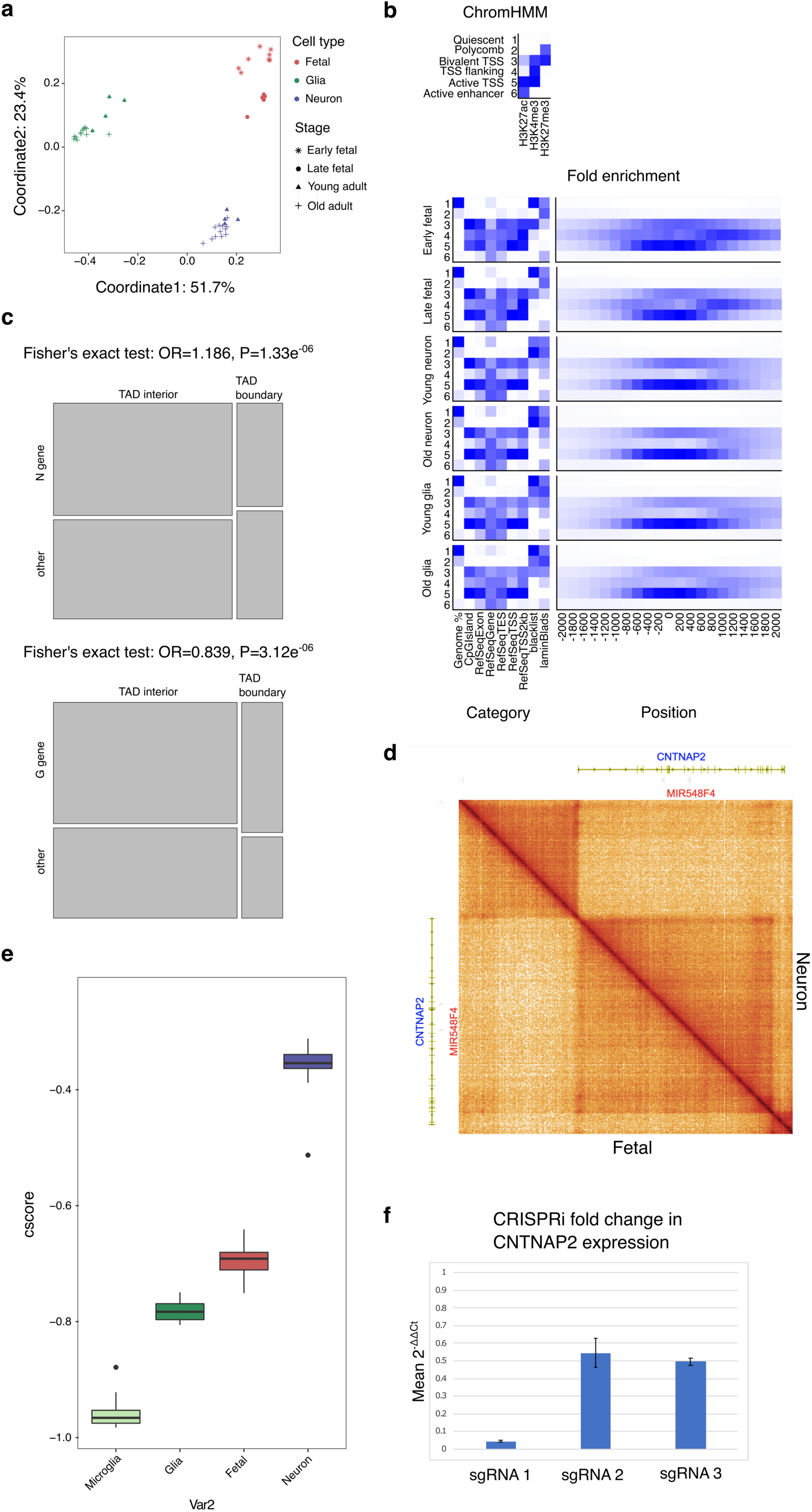
**a**, Multidimensional scaling plot showing differential TADs between different cell types and developmental stages (% variance explained). **b**, Functional chromatin states defined by ChromHMM and their fold enrichment across different categories of regulatory sequences and across positions in proximity to transcription start sites of genes. **c**, Fisher’s exact test to determine likelihood of neuronal differential gene expression at neuronal differential TAD boundaries (top) and likelihood of glial differential gene expression at glial differential TAD boundaries (bottom). OR: odds ratio, P: P-value. **d**, Snapshot of the *CNTNAP2* locus from the HiGlass 3D genome browser showing Hi-C contact maps (10 kb resolution) in fetal and adult neurons. **e**, Mean cscore and variance −50kb to +500 kb from TSS of *CNTNAP2* in different cell types. Pairwise differences between cell types are statistically significant (p-value=3.05e^−05^). **f**, Fold change in *CNTNAP2* expression upon CRISPRi. Mean 2^−**ΔΔCt**^ was determined across 3 replicates relative to the scrambled guide control, normalized over the geometric mean of housekeeping genes *GAPDH* and *ACTB*. sgRNA 1: mean 2^−**ΔΔCt**^ **=** 0.044497 (SEM= 0.0046653), sgRNA 2: mean 2^−**ΔΔCt**^ **=** 0.54475 (SEM= 0.080944), sgRNA 3: mean 2^−**ΔΔCt**^ **=** 0.49711 (SEM= 0.020382). Hi-C data was generated from cells transduced with sgRNA 1, the top scoring guide from the Broad Institute CRISPRi algorithm.

**Supplementary Fig. 3.**
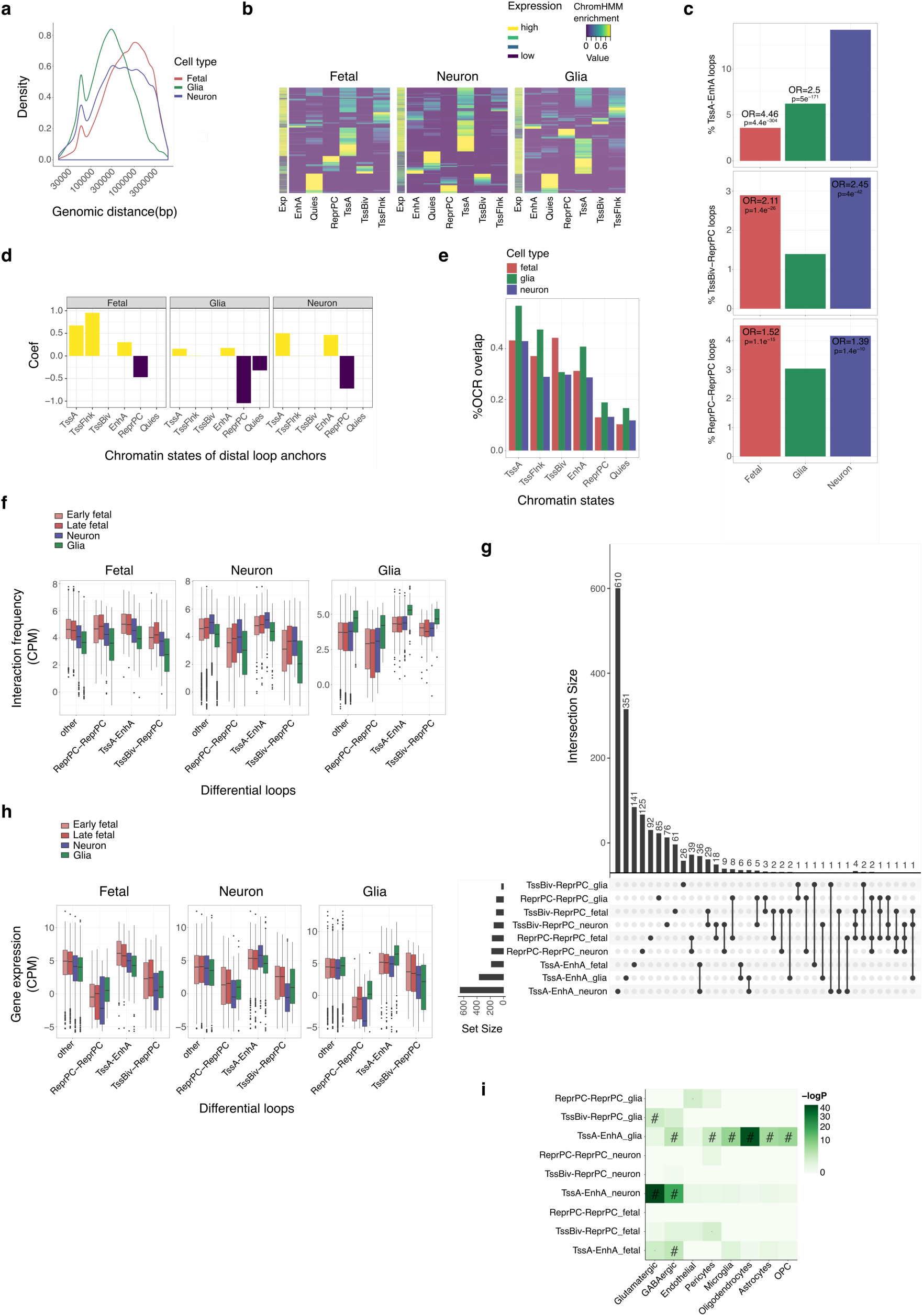
**a**, Frequency distributions of loop sizes across merged Hi-C data from fetal neurons, adult neurons, and glia. **b**, Chromatin states (% enrichment) at loop associated promoters [-1 to +1 kb from TSS] and gene expression levels. TssA: active transcription start site (H3K4me3, H3K27ac), TssFlnk: transcription start site flanking (H3K4me3), TssBiv: bivalent transcription start site (H3K4me3, H3K27me3), EnhA: active enhancer (H3K27ac), ReprPC: polycomb repressed (H3K27me3), Quies: Quiescent. **c**, Proportion of loops classified as TssA-EnhA, TssBiv-ReprPC, or ReprPC-ReprPC. TssA-EnhA frequency in adult neurons expressed as odds ratio relative to fetal neurons and glia. TssBiv-ReprPC and ReprPC-ReprPC frequencies in fetal and adult neurons expressed as odds ratio relative to glia. **d**, Prediction coefficients of gene expression change from the chromatin state of the distal loop anchor using Lasso regression. Fetal spearman correlation=0.29, Neuron spearman correlation=0.34, Glia spearman correlation=0.31. **e**, Percent of open chromatin regions (OCRs) obtained from ATAC-seq that overlap with chromatin state assigned loop anchors. **f**, Interaction frequencies (counts per million reads mapped) of differential promoter-anchored loops across cell types and fetal developmental stages. **g**, UpsetR plot showing intersections between differential promoter-anchored loops. **h**, Gene expression levels (counts per million reads mapped) associated with differential promoter-anchored loops across cell types and fetal developmental stages. **i**, Cell type specificity of chromatin state assigned differential promoter-anchored loops.

**Supplementary Fig. 4.**
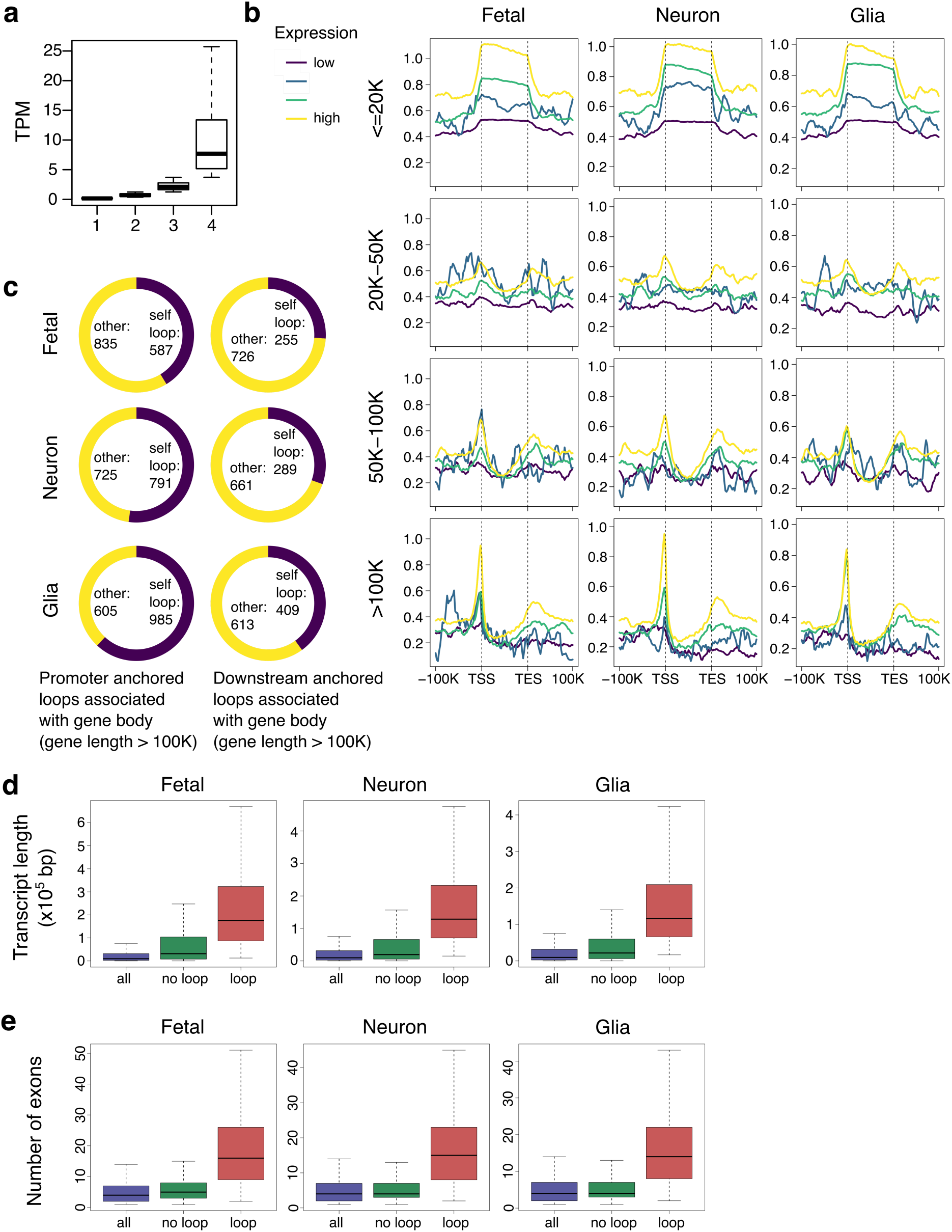
**a**, TPM (transcripts per million reads mapped) distributions across 4 expression levels. **b**, Loop anchor densities from TSS (transcription start site) to TES (transcription end site) across 4 different expression levels for different gene length categories (<=20 kb, 20-50 kb, 50-100 kb, >100 kb). **c**, Percent of promoter-anchored loops and percent of downstream anchored loops that are associated with the gene body (self-loops). **d**, Transcript length distributions in all genes, genes with no exon-exon loops, and genes with exon-exon loops. **e**, Exon number distributions in all genes, genes with no exon-exon loops, and genes with exon-exon loops.

## Supplementary Tables

**Supplementary Table 1**: Processing of paired-end reads from replicate Hi-C libraries. Excel sheet attached separately.

**Supplementary Table 2a.**
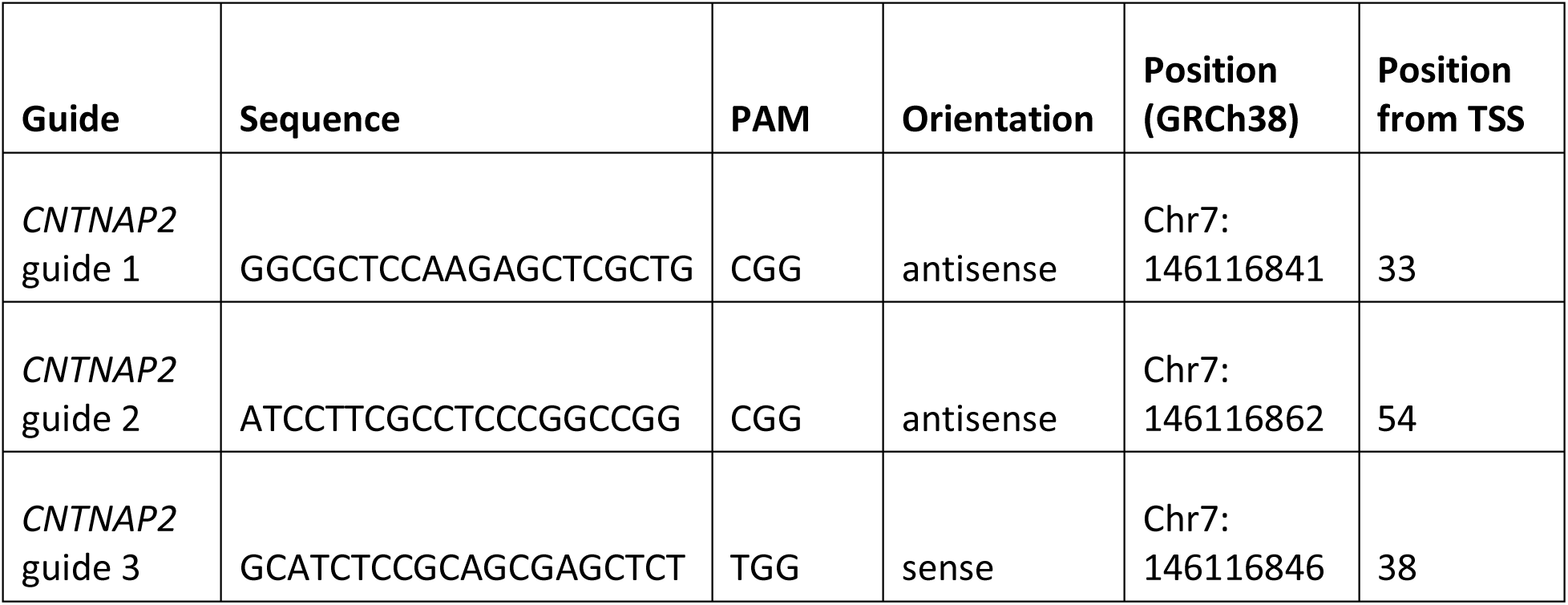
*CNTNAP2* CRISPRi guides designed with the BROAD Institute CRISPRi algorithm and validated *in vitro*.

**Supplementary Table 2b.** TaqMan probe used for CRISPRi validation.

| Gene | TaqMan Assay ID | Chromosomal location (GRCh38) | Amplicon size (bp) | Probe spans exons |
| --- | --- | --- | --- | --- |
| *CNTNAP2* | Hs01034296_m1 | Chr.7:<br>146116035 -<br>148420998 | 63 | Yes |

**Supplementary Table 3a:** Spatial relationship between chromatin state assigned promoter-anchored loops and TADs in each cell type. Excel sheet attached separately.

**Supplementary Table 3b:** Overlap between enhancer-promoter loops and differential TADs in adult neurons and glia.

|  | no overlap with diffTAD | Promoter loop anchor overlaps with diffTAD | Distal enhancer loop anchor overlaps with diffTAD | Both loop anchors overlap with diffTAD |
| --- | --- | --- | --- | --- |
| **Neuron** | 2759 | 67 | 5 | 2 |
| **Glia** | 1439 | 6 | 1 | 0 |

|  | Neuron | Glia |
| --- | --- | --- |
| **E-P loop anchors overlap with diffTAD** | 74 | 7 |
| **no overlap with diffTAD** | 2759 | 1439 |

**Odds ratio of Neuron vs Glia:** 5.51289
**p value** 1.54E<sup>-07</sup>

**Supplementary Table 3c:** Biological processes associated with genes that are enriched for E-P loops that overlap with neuronal differential TADs. GO terms were obtained with g:Profiler ^88^, which uses multiple testing correction to discard false positives.

| Biological processes | -log10_of_adjusted_p_value |
| --- | --- |
| synapse assembly | 8.220835447 |
| synapse organization | 5.030304365 |
| cell junction assembly | 4.927943433 |
| synaptic membrane adhesion | 3.826264391 |
| regulation of synapse assembly | 3.184423975 |
| cell junction organization | 3.120539228 |
| presynapse assembly | 2.807211942 |
| presynapse organization | 2.710079666 |
| positive regulation of synapse assembly | 2.448280578 |
| regulation of cell junction assembly | 1.753378469 |
| regulation of synapse organization | 1.655147558 |
| positive regulation of cell junction assembly | 1.63280908 |
| regulation of synapse structure or activity | 1.552411177 |

**Supplementary Table 3d**: Genes enriched for E-P loops that overlap with neuronal differential TADs. Excel sheet attached separately.

**Supplementary Table 3e**: Genes associated with differential chromatin state assigned promoter-anchored loops in each cell type. Excel sheet attached separately.

## Notes

### Competing Interest Statement

The authors have declared no competing interest.

